# Transcriptomic Analysis Reveals Inflammatory and Metabolic Dysregulation in Unexplained Female Infertility

**DOI:** 10.64898/2026.01.24.701467

**Authors:** Ritika Patial, Sonalika Ray, Kashmir Singh, R.C. Sobti

**Affiliations:** Centre for Systems Biology and Bioinformatics, Panjab University, Chandigarh; Research Consultant, Louisiana State University, Baton Rouge, USA; Department of Biotechnology, Panjab University, Chandigarh

**Keywords:** Infertility, Gene expression data analysis, Differential gene expression analysis, KEGG pathways, Biological processes, Molecular Functions, Cellular Components, Protein-protein interaction (PPI) network, WGCNA, IPA (Ingenuity Pathway Analysis), Endometrial tissue

## Abstract

Infertility is a complex condition affecting both the male and female population. Influenced by multiple factors, it remains a constant challenge due to limited understanding of endometrial abnormalities. With this study we aim to investigate the molecular basis of infertility using transcriptomic analysis of endometrial tissue from the NCBI GEO dataset GSE92324. We performed exploratory data analysis using Principal Component Analysis (PCA) to find samples variance followed by differential gene expression (DGE) analysis using DESeq2 package where we identified 168 significant genes with adjusted p-value < 0.05 and |log2FC| > 2. Upregulated genes included *GPX3, CXCL14,* and *PPARGC1A* and downregulated genes included *WNK4, GJB2*, and *TRPM6*. Functional enrichment using KEGG and GO showed that differentially expressed genes (DEGs) are involved in immune-inflammatory pathways, lipid metabolism and steroid biosynthesis pathways. Through Ingenuity Pathway Analysis (IPA) we identified affected canonical pathways such as increased innate immune responses, altered lipid metabolism and inhibition of mitochondrial dysfunction. Upstream regulator analysis highlighted PTEN, PRKAA1, HDAC4, IL10RA, and RAD51, which were impacting metabolic pathways and anti-inflammatory signalling. Further, through Weighted Gene Co-expression Network Analysis (WGCNA) we found a Turquoise module that had very strong and highly significant negative correlation (cor = - 0.84, respectively and P < 0.0001) with traits of interest. This led to the discovery of C7orf50 as a novel insight involved in cholesterol metabolism linked to infertility. This integrative approach reveals crucial genes, co-expression modules, and underlying pathways involved in female infertility.

**GRAPHICAL ABSTRACT:** 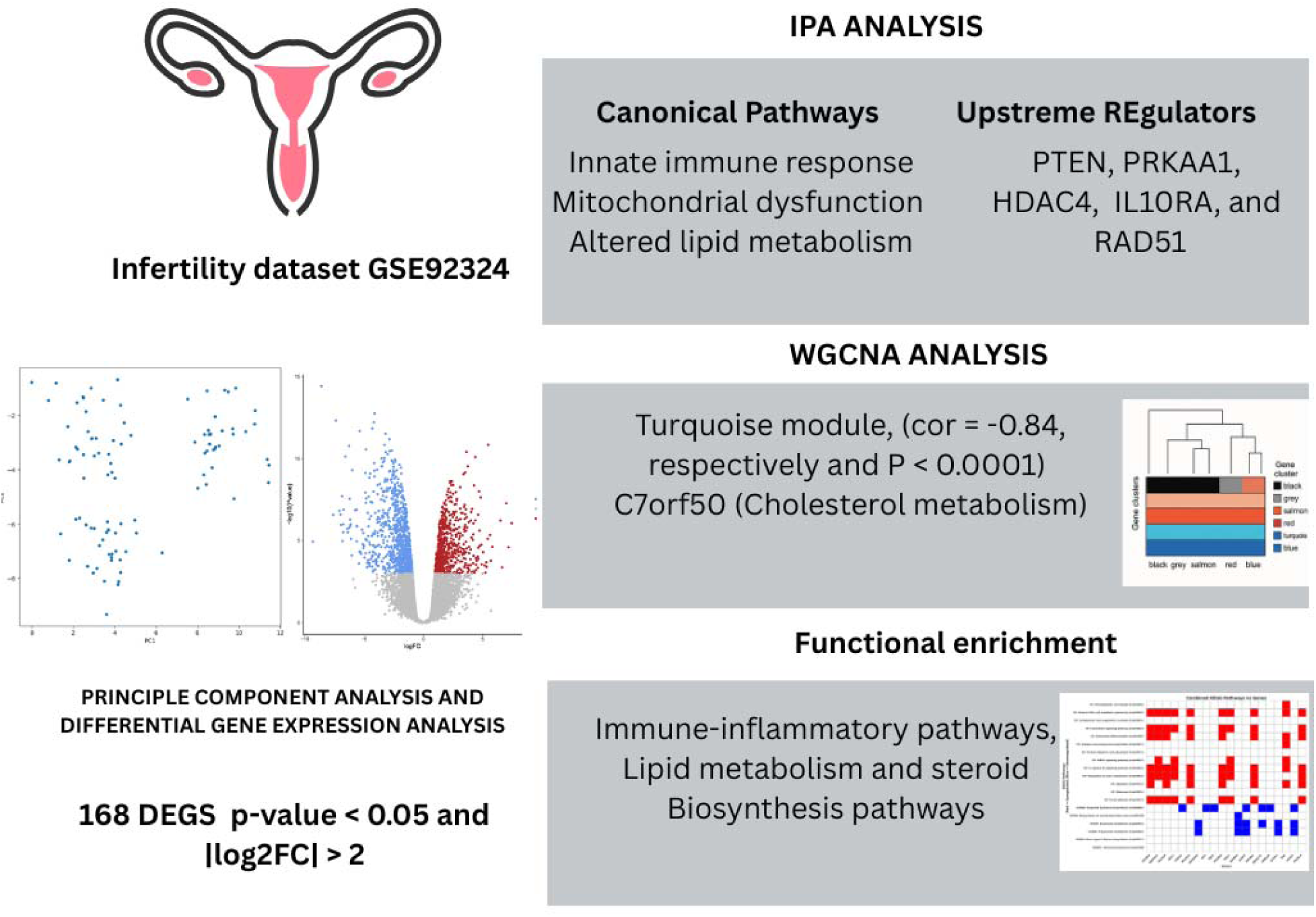

**HIGHLIGHTS:**

- From the dataset GSE92324 total of 168 significant DEGs associated with unexplained infertility were identified using adjusted p-value < 0.05 and |log2FC| > and < 2.
- In comparison with the CTD list we identified five genes C1orf106, C15orf59, LINC00461, C15orf48, and C10orf99 previously unknown as having direct evidence of involvement in infertility.
- WGCNA analysis highlighted the turquoise module as highly associated and gave the novel gene C7orf50 associated with cholesterol metabolism.
- IPA revealed PTEN, PRKAA1, IL10RA, and RAD51 as potential upstream regulators and inflammatory pathways, mitochondrial dysfunction as canonical pathways.
- The study highlights a novel link between GI inflammation and endometrial receptivity.

## 1. INTRODUCTION

Infertility is a significant health issue affecting a large proportion of the female population, with numerous underlying causes from hormonal imbalances to structural abnormalities. The global prevalence of infertility is estimated to be around 12.87 percent [1], with an average annual increase of 0.68 from 1990-2021 [2]. According to recent statistics, one in every six people of reproductive age worldwide will experience infertility in their lifetime . In 2021, this translated to roughly 110 million women (3.7% of women aged 15-49) and 55 million men (1.8% of men aged 15-49) living with infertility globally [2]. It has been observed that over a decade the prevalence of infertility has been increasing. This rise is attributed to multiple factors, including demographic changes and lifestyle shifts. Hormonal imbalances are considered as the root cause of female infertility, as it affects ovulation and menstrual cycles [3]. It is important to have proper endocrine function as it is essential for ovulation and reproductive cyclicity. Disorders of the *hypothalamic-pituitary-ovarian (HPO) axis* or other endocrine glands can disrupt the delicate hormonal balance required for follicle development, ovulation, and implantation [4]. Abnormalities in fallopian tubes, their blockage, uterine fibroids and endometriosis are also amongst the anomalies that also impede the implantation process [5]. For infertility genetic factors are also contributing, conditions like ovarian insufficiency, mullerian anomalies have the capabilities to affect reproductive function [6]. The genetic factors may range from chromosomal disorders to single-gene mutations and polymorphisms that affect reproductive function. A well-known example is the *FMR1* gene premutation (associated with Fragile X syndrome carriers), which confers about a 20% risk of developing premature ovarian insufficiency - a tenfold higher risk than in the general population [7]. This leads to depletion of ovarian follicles before age 40, causing infertility. Other genetic contributors include mutations in genes critical for oocyte meiosis, ovarian development, or hormone signaling. Changing lifestyle is also one of the dominating factors in case of infertility. The nutrition of the individual, either being under nutrient or overweight also impacts female infertility. Women who are significantly underweight or overweight often experience disrupted ovulation and menstrual irregularities. High adiposity, for example, alters peripheral estrogen and insulin levels, contributing to anovulation (as seen in many cases of PCOS). Frequent consumption of trans-fats and ultra-processed foods has been associated with ovulatory infertility, whereas diets rich in unsaturated fats, whole grains, and antioxidants are linked to better fertility outcomes [8]. Stress and depression can also likewise affect fertility directly through elevated cortisol levels or through adoption of unhealthy habits. Additional substance abuse such as smoking which causes ovarian follicle disruption, alcohol intake harms oocyte quality and disrupts menstrual cycle with poor sleep increasing the negative impact on fertility [9]. Poor sleep, due to personal choices, late working hours, more exposure to artificial light alters the circadian cycle of the individuals. Studies have established a strong link between circadian cycle and hormone signalling in the human body. Disruption of these rhythms dysregulation of the hypothalamic-pituitary-gonadal axis, causing irregular secretion of GnRH, LH, and FSH which in return affect ovulation and luteal function impacting ovarian function and endometrial receptivity [10]. With its impact over a significant portion of the population and the level of complexity it has due to genetic, epigenetic and also the lifestyle and environment factors, understanding the molecular mechanisms behind infertility has become very crucial for developing effective diagnostic and therapeutic strategies. With advancements in the field of genetics and molecular biology deeper insights have been gained but still there are few factors that are uncategorized. While advances in genetics and molecular biology have provided deeper insights, some factors remain uncategorized. The advancement in bioinformatics tools have expanded our understanding of gene expression profiling in infertility. Dong et al., 2022 have investigated the association of cell cycle, immune response as targets for addressing infertility [11]. A study by Luz et al., 2022 also highlighted the role of cytokine signalling and inflammatory responses [12]. Studies by Vargas et al., 2023 and Bui et al., 2024 highlighted pathways affecting endometrial receptivity which also included immune response and inflammatory responses [13,14]. Despite these insights and advancements, there remains a lack of extensive analysis into biomarkers and pathways and their biological significance as these studies only focus on immune responses and cytokine responses. Infertility is now becoming a lifestyle based disorder, influenced by various modifiable factors that affect female reproductive health. While genetic and medical conditions play a role, lifestyle choices such as diet, physical activity, smoking, alcohol consumption and higher stress levels have been shown to significantly impact fertility [8,15]. Therefore there is a significant gap into these domains and they should be explored. Therefore the study aims to address this gap by using a bioinformatics based approach to identify DEGs and perform their functional enrichment to understand their biological significance. We further employ Weighted Gene Co-expression Network Analysis (WGCNA) to explore gene co-expression patterns, and perform IPA to identify significantly affected canonical pathways, disease and biofunctions, as well as ML predicted disease pathways, providing a higher-confidence understanding of the molecular mechanisms involved in infertility.

## 2. METHODOLOGY

### 2.1 Data Extraction

Gene expression data for this study was obtained from the NCBI-GEO (GSE92324) [16], which includes transcriptomic profiles of endometrial tissue collected during the implantation window. The dataset consists of samples from two groups: cases of implantation failure in women with unexplained infertility and proven fertile oocyte donors serving as controls. A total of 18 samples were included: 10 from implantation failure cases and 8 from healthy controls, all collected under the influence of ovarian stimulation during an IVF cycle. After retrieval, the raw data was preprocessed to ensure integrity and comparability. In the preprocessing step the duplicate gene symbols, genes with zero or very low expression values, and genes with no measurable variance across samples were removed so that only significant genes with measurable differences in expression between cases and controls were retained for downstream analysis.

### 2.2 Data Normalization

Following the pre-processing step, the next step was to normalize the expression matrix using log transformation as it stabilizes variance, minimizes the influence of extreme expression values, and provides normal distribution across the data [17]. Normalization is a crucial preprocessing step in RNA-Seq workflows, especially in differential expression analysis or in PCA where dimensionality reduction is needed [18]. It makes sure that technical artifacts and variability don’t hide real biological differences, which makes downstream statistical models more accurate and easier to understand.

### 2.3 Exploratory Data Analysis

PCA on the log-transformed data was done, to assess the variability in the dataset and to look for any potential outliers. PCA is a dimensionality reduction technique that reduces the dimensionality of the data to make visualization of variation of points in the data easier [19]. The data was log-transformed prior to PCA to ensure that genes with higher expression levels do not have undue influence on the principal components.

### 2.4 Differential Gene Expression Analysis

DGE analysis between control and case samples using the DESeq2 package was done [20]. To focus on the most statistically significant DEGs, stringent filtering criteria was applied. The Genes were considered statistically significant if they had p-value < 0.05 and an adjusted p-value (padj) < 0.05 [21]. Also to focus on more biologically relevant genes the filter on Log2 fold change of ≥ 2 or ≤ -2 was applied. Further, to visually represent the results, we used the EnhancedVolcano package in R to create a volcano plot, which highlights genes based on both statistical significance and magnitude of expression change.

### 2.3 Functional Characterization

To understand the biological impact of the DEGs, functional enrichment analysis using the ClusterProfiler package in R was done [22], following DGE analysis. This package enables statistical enrichment analysis and visualization. Also, to understand the involvement in biological processes (BP), molecular functions (MF), and cellular components (CC) associated with the DEGs [23], Gene Ontology (GO) enrichment was performed. In addition, KEGG pathway enrichment analysis was performed to understand the involvement of DEGs in known signaling and metabolic pathways [24].

### 2.4 Ingenuity Pathway Analysis (IPA)

To complement the R based pathway analysis, IPA was done to gain deeper biological insights from DEGs [25]. Canonical pathways using a threshold of < 0.05 for p value, z-scores of ≥ 2 and ≤ -2 for activated and inhibited pathways were applied [26]. The disease and function analysis defines DEGs into different biological groups such as inflammation, reproductive dysfunction, or hormone regulation, explaining how dysregulation in genes may contribute to phenotype. In the upstream regulator analysis, transcription factors, cytokines, and signaling molecules potentially responsible for the observed gene expression changes are identified with the filter of z-scores ≥ 2 or ≤ -2 and significant p value.

### 2.5 Weighted Gene Co-Expression Network Analysis (WGCNA)

The R package WGCNA was used to perform the WGCNA analysis on gene expression dataset [27]. Initially, a cluster analysis was carried out to check for any potential outliers within the samples. Subsequently, to prevent the arbitrary selection of a cut-off point, an optimal soft threshold power β for network construction was determined using the Pick Soft Threshold function within the ’WGCNA’ package. After the identification of the soft thresholding power β, to enable the weighted separation of co-expression patterns, an adjacency matrix was created. By using Topological Overlap Measure (TOM) which is a network distance measure, the co-expression similarity between each pair of genes from the adjacency matrix was calculated. The utilization of TOM helped in reducing the impact of noise and unwanted associations. Finally, the co-expression gene modules were detected through hierarchical cluster analysis, and modules showing a high level of similarity with a clustering height of module eigengene (ME) lower than 0.25 were merged to streamline the analysis.

### 2.6 Module-Trait Relationship

The relationships between co-expression gene modules and the disease was analysed using the Pearson Correlation Coefficient (PCC) for each module and the specific clinical trait under investigation. Subsequently, a t-test was utilized to determine the p-value, with a threshold of <0.05 for being considered statistically significant. The emphasis was on modules highly linked to the trait, and the genes within these modules were identified through the computation of Gene Significance (GS) and Module Membership (MM). Gene Significance (GS) was defined as the absolute correlation between a gene and a specific clinical parameter, while Module Membership (MM) indicated the correlation between the module eigengene (ME) and the gene’s expression pattern. Genes exhibiting high values for both GS and MM were classified as hub genes within a module.

### 2.7 Systems Biology Approach and Network Construction

To gain deeper insight into the molecular interactions among the DEGs, a systems biology approach using protein-protein interaction (PPI) network analysis was done. The list of DEGs (fold change of ≥ 2 or ≤ -2 and padj < 0.05) was submitted to the STRING database to retrieve high-confidence interaction data. The threshold was set at a confidence level of 0.700 to lower the possibility of false positives and focus on interactions that are important to biology. Then, the network was imported into Cytoscape, to visualize and analyze biological networks. The hub gene analysis was performed using the CYTOHUBBA plugin, in Cytoscape.

## 3. Results

### 3.1 Exploratory Analysis

In the PCA plot (Figure 1), the control group, shown within the blue ellipse, formed a tight cluster, reflecting low variability and consistent gene expression profiles across these samples. On the other hand, the infertility group (red ellipse) formed a distinct but the samples were more dispersed in the cluster, indicating higher heterogeneity among these samples **Figure 1**. The first principal component (PC1) captures 46.62% of the variance, while the second principal component (PC2) captures an additional 10.97%, highlighting strong transcriptional differences related to implantation failure. Also, there was no identification of outliers in the respective groups.

**Figure 1:**
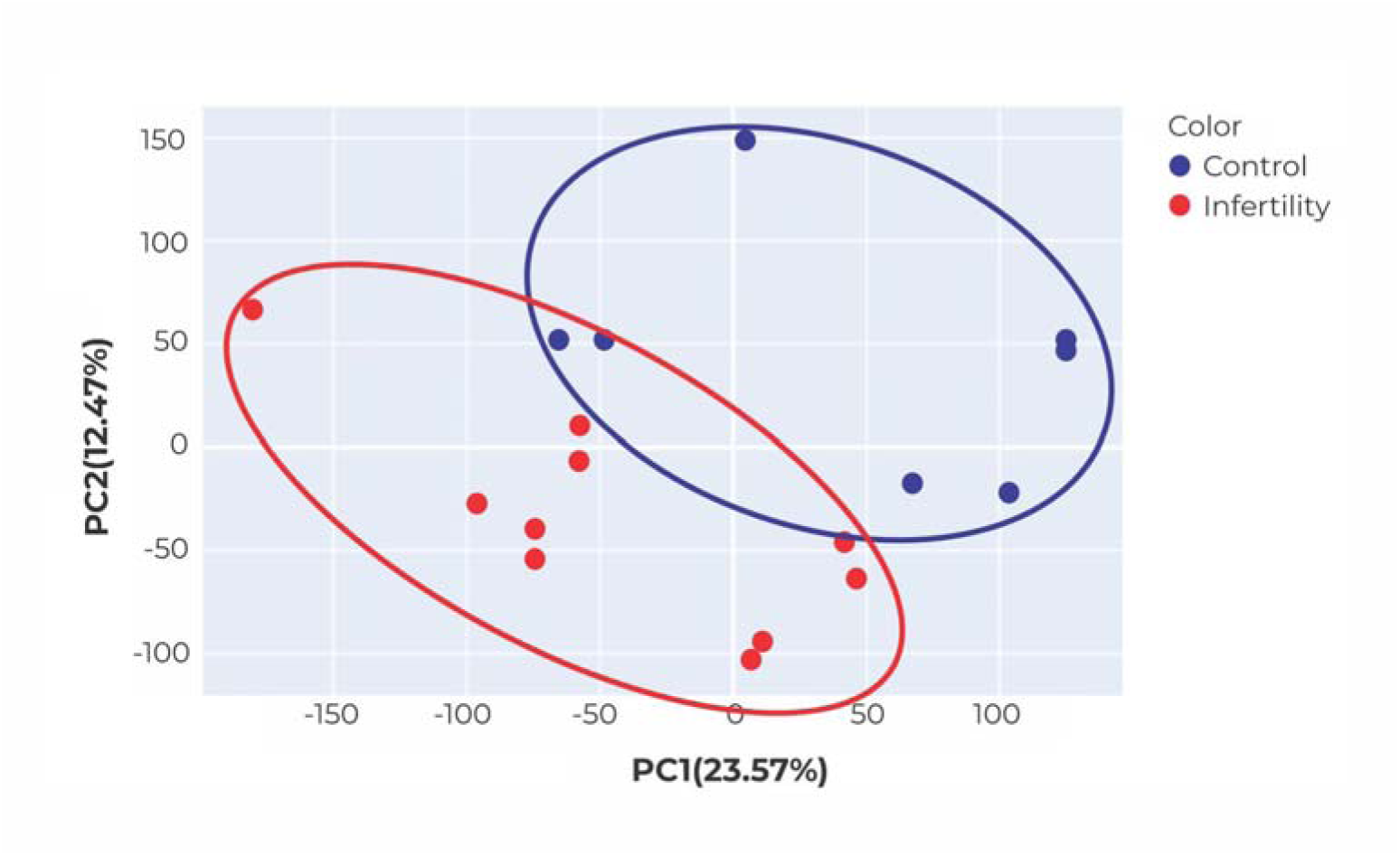
Principal Component Analysis (PCA) of the dataset. The blue ellipse represents control samples, and the red ellipse represents samples from unexplained infertility cases.

### 3.2 Differential Gene Expression (DGE) Analysis

A comprehensive DGE analysis using DESeq2 between infertility and control samples was done. A total of 1,775 significant DEGs were initially identified at a p.adj < 0.05. To further emphasize on the biologically significant findings, stringent filtering was applied of absolute log fold change (|log FC|) > 2, which resulted in 168 DEGs identification. Out of total 47 were upregulated and 121 were downregulated genes. Among the top significantly upregulated genes, GPX3 (log FC = 4.62), CXCL14 (4.40), AOX1 (3.91), HSFX1 (3.75), CDA (3.48), PPARGC1A (3.25), TSPAN8 (3.24), NNMT (3.10), LAMB3 (3.09), and C3 (3.01) were notably enriched. These genes have well known association in context of infertility pathophysiology, associated with oxidative stress response (GPX3 [28], inflammation and immune modulation (CXCL14, C3), metabolic reprogramming (AOX1, NNMT [29,30]), mitochondrial biogenesis and energy metabolism (PPARGC1A [31]), and extracellular matrix remodeling (TSPAN8, LAMB3 [7,8]).

Also, the most significantly downregulated genes included WNK4 (4.20), GJB2 ( 4.25), TRPM6 (4.27), CAPN6 (4.42), HPCAL4 (4.60), OPRPN (4.67), CHST4 (4.70), GP2 (5.07), NLRP5 (5.17), and SLC7A4 (5.54). These genes are associated with pathways that are crucial in fertility processes such as ion transport and cellular signaling (WNK4, TRPM6 [32,33], cell-cell communication and gap junctional intercellular coupling (GJB2 [34], proteolysis and cellular regulation (CAPN6 [35], immune and inflammatory response modulation (OPRPN, GP2, CHST4), maternal effect and embryo development (NLRP5 [36,37]), and amino acid transport (SLC7A4 [38]. The volcano plot (**Figure 2**) represents upregulated and downregulated genes with significant log fold change.

**Figure 2:**
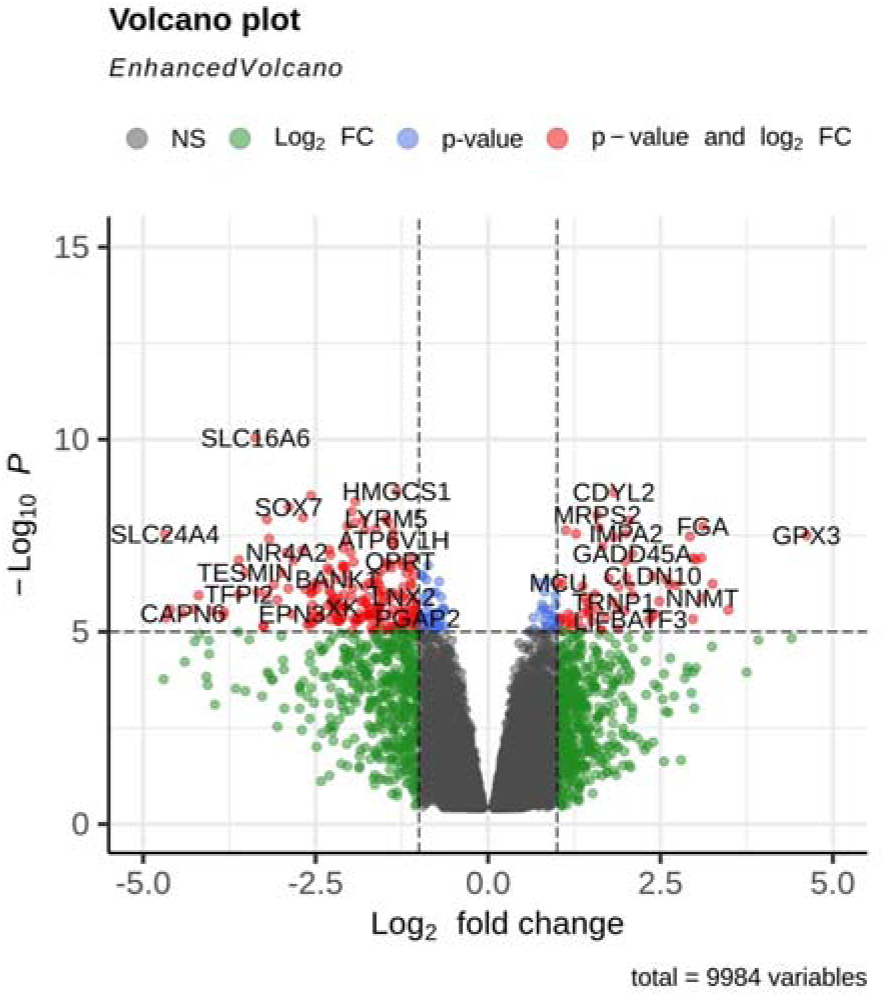
Volcano plot showing the DEGs between control and infertility cases. Genes with significant changes are highlighted.

Comparison of these DEGs with the CTD highlighted that none of the 168 DEGs have known curated therapeutic or marker/mechanism roles directly associated with female infertility. However, 163 of these DEGs had hypothetical or theoretical associations according to CTD. Additionally, literature mining identified five genes C1orf106, C15orf59, LINC00461, C15orf48, and C10orf99, previously unknown as having direct evidence of involvement in infertility. These genes thus present novel opportunities for infertility research.

### 3.3 Co-expression Network Analysis (WGCNA)

#### Hierarchical Clustering and Module Detection

We assessed the quality of the data by clustering the samples based on the distance between various samples found in Pearson’s correlation matrices and creating a dendrogram. The clusters did not contain any outliers, thus a total of 18 samples were further used in the analysis. Subsequently, a power value of β = 10 was chosen as the soft threshold to ensure that there is a scale-free network (**Figure 3 C**). Consequently, a total of 45 modules were formed by grouping 8245 genes using the average linkage hierarchical clustering algorithm. Further we established the correlation between each module and all the clinical information (condition) in the GSE92324 dataset by computing the MS for each module-trait correlation. From the analysis we discovered the Turquoise module showed very strong and highly significant negative correlation (cor = -0.84, respectively and P < 0.0001) and purple module showed positive correlation (cor = +0.65, respectively and P < 0.001) but slightly weaker than turquoise **Figure 3 A**. Further to gain deeper insights we compared the genes of the module against the (DEG) list. The comparison showed that there were many significant numbers of genes that are common between the DEGs and the module-associated genes, suggesting possible functional relevance and so supporting the function of these genes in the basic biological condition. Through literature reading, it was found that C7orf50, LOC652276, and CASTOR3 are novel insights in the WGCNA module. LOC652276, and CASTOR3 are the pseudogenes with no relevance to infertility. Whereas, C7orf50 is involved in cholesterol metabolism, but its context to female infertility is unknown **[39].**

**Figure 3:**
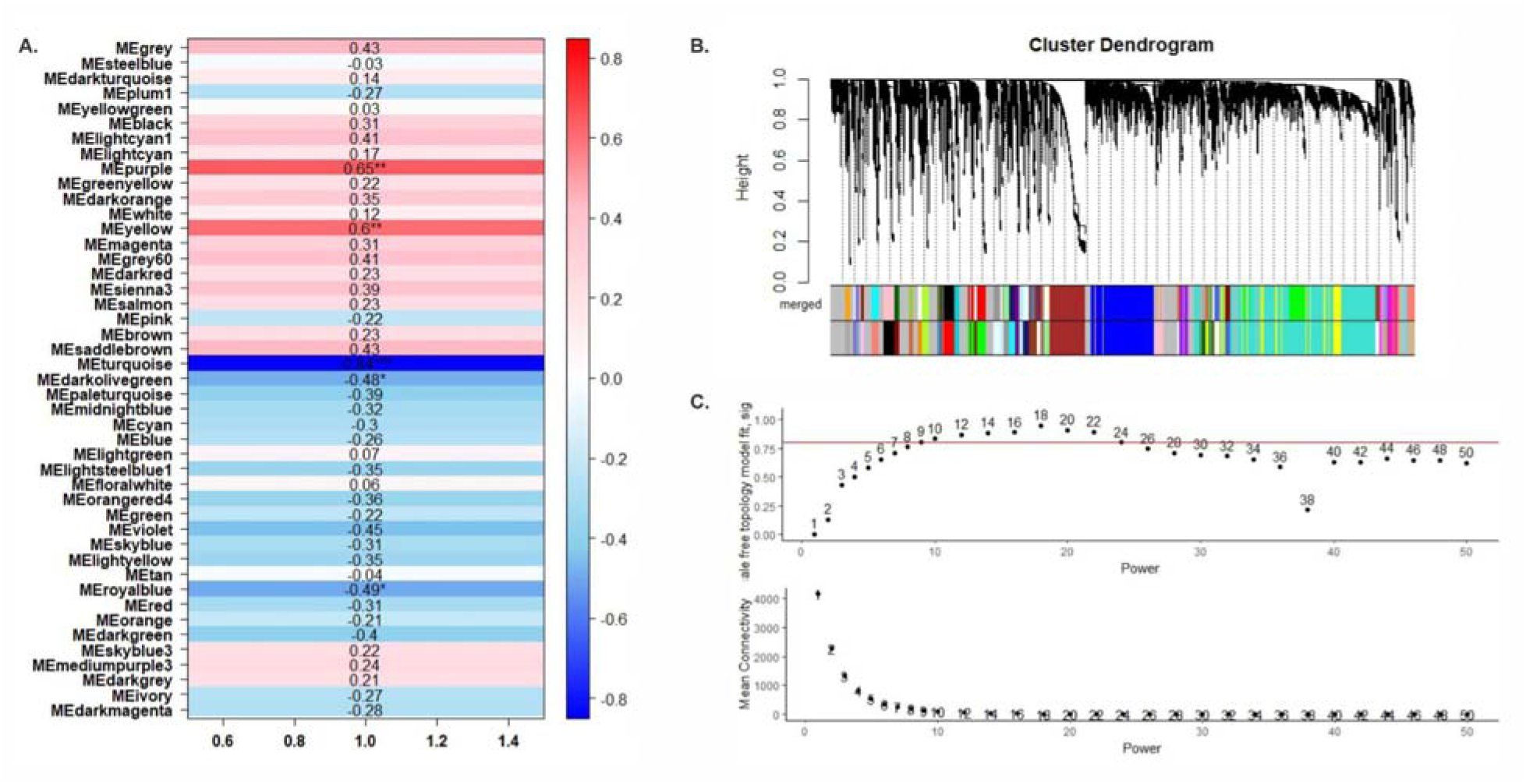
WGCNA analysis of transcriptomic data. A. Heatmap correlating module eigengenes with traits of interest. Color represent strength of correlation (red: positive, blue: negative), with asterisks indicating statistical significance (*p < 0.05, **p < 0.01, ***p < 0.001). B. Cluster dendrogram showing hierarchical clustering of gene expression profiles, with dynamic tree cut used to identify modules. Color bands represent merged modules. Scale-free topology fit index and mean connectivity analysis to determine the optimal soft-thresholding power for network construction. Power = 10 achieves high scale-free topology with acceptable mean connectivity.

### 3.4 Functional Enrichment Analysis

#### 3.4.2 KEGG Pathway Enrichment

Functional enrichment analysis using the KEGG identified several significantly upregulated pathways as shown in Figure 4 and Table 1. It included Hematopoietic Cell Lineage (p = 0.0005), Natural Killer Cell Mediated Cytotoxicity (p = 0.0008), and Complement and Coagulation Cascades (p = 0.0014) **Figure 4A**. Some of the genes that were more common in these pathways were CR1, CSF1R, TNF, CD3E, and members of the PIK3 family. NK cells are abundant in the endometrium (as uterine NK cells) and play complex roles in implantation such as facilitating trophoblast invasion and vascular remodeling in a restrained manner. However, an enrichment of *cytotoxicity* pathways suggests an excessive or over-active NK cell phenotype in the infertile endometrium. This is in line with clinical observations that dysregulated NK cell activity can be harmful, for example, elevated NK cell cytotoxicity (and not just higher NK cell numbers) has been implicated in recurrent pregnancy loss and unexplained infertility **[40]**. Uterine NK cells in these patients are overly cytotoxic, they might attack the implanting embryo or disturb the tolerance normally required at the maternal-fetal interface **[41]**.

**Table 1:**
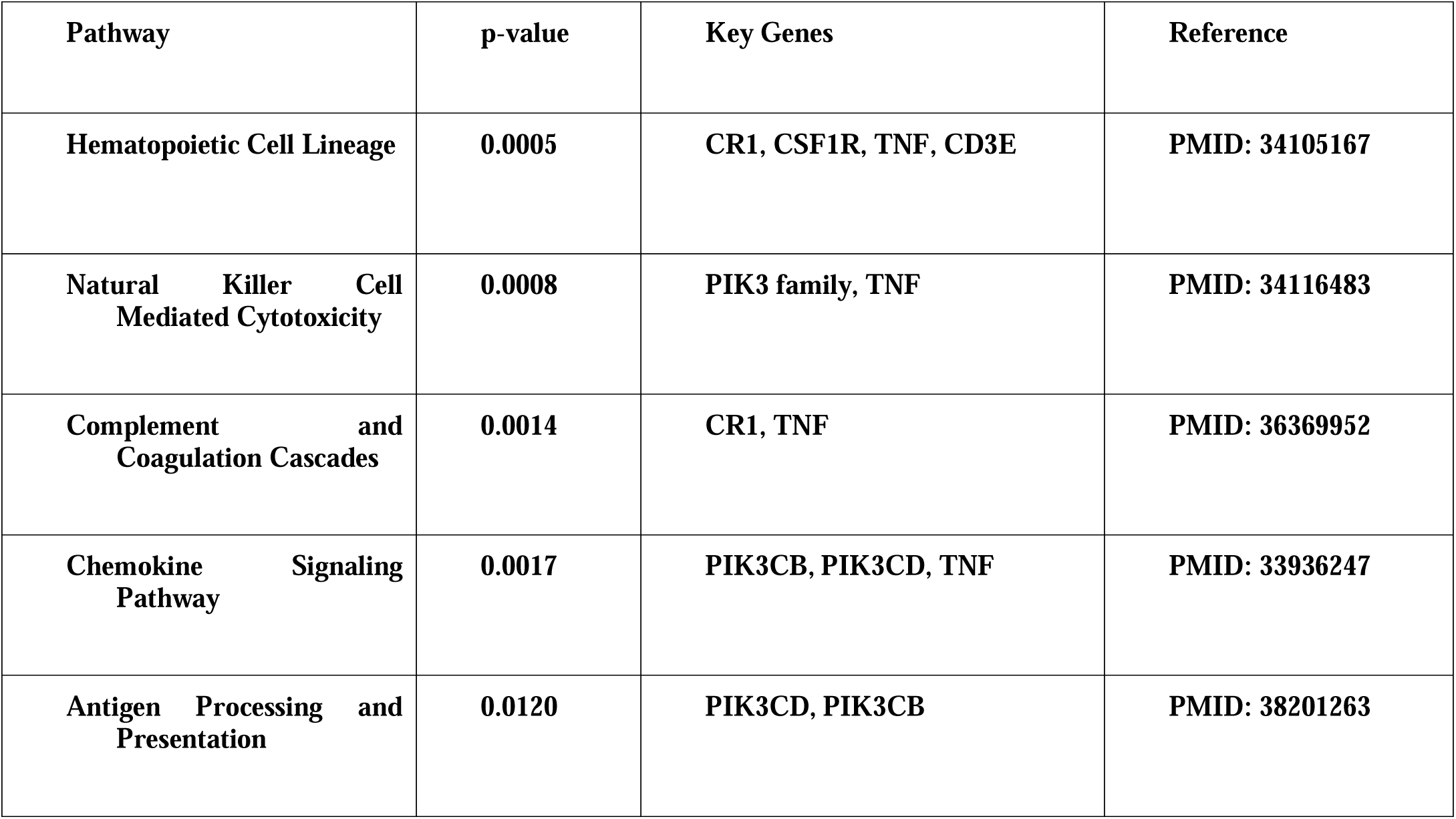
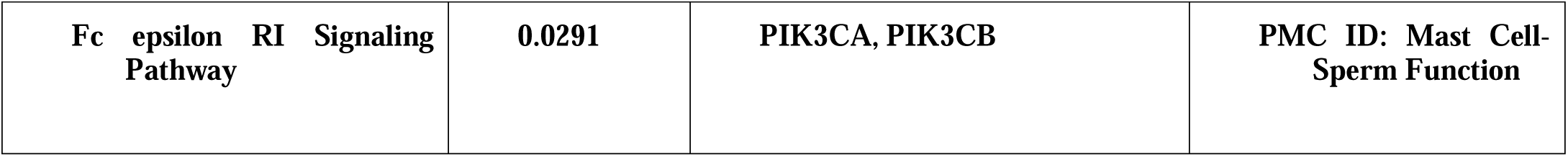
Significantly Upregulated KEGG Pathways.

**Table 2:**
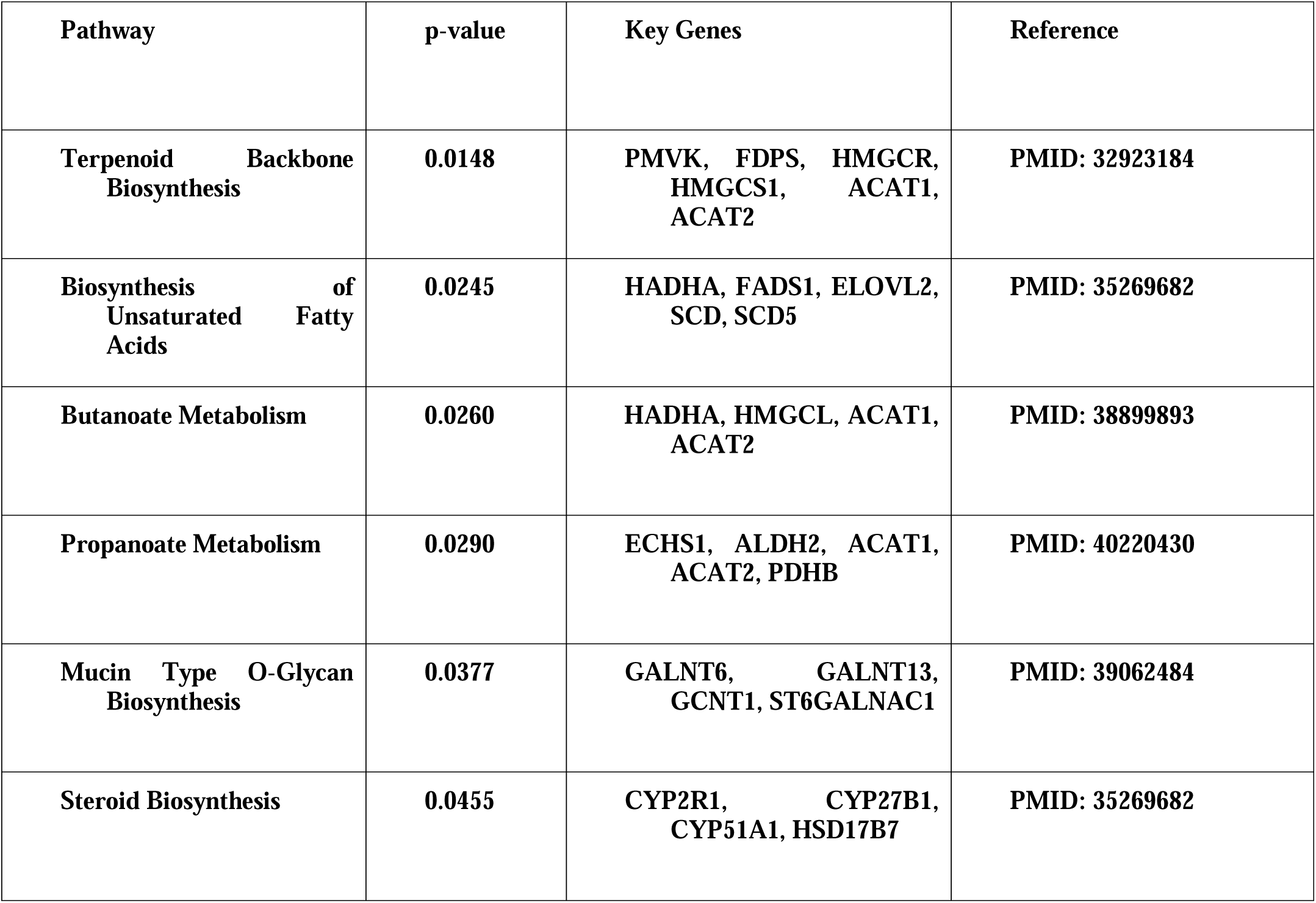
Significantly Downregulated KEGG Pathways.

Alongside NK cells, the complement cascade appears upregulated. Components of the complement system (C1, C3, C4, etc.) were among the upregulated genes, suggesting an active complement system in the endometrium. During normal early pregnancy the activity of complement at the site of implantation is strictly regulated, while remodeling [42,43]. But its over-activation must also be stopped to prevent tissue damage to the developing embryo. An unchecked complement cascade in the absence of an embryo could indicate a pro-inflammatory microenvironment. Complement proteins can promote recruitment of immune cells and release of inflammatory mediators that might degrade the endometrial tissue or vasculature. As a matter of fact, excessive complement activation has been linked to pregnancy complications (e.g., mutations in complement regulatory proteins are associated with preeclampsia), underscoring that the complement system at the implantation site can be a double-edged sword [44]. Therefore, the enrichment of complement pathways in our analysis further supports the idea of an inflammation-rich endometrial environment in unexplained infertility. Additional pathways such as the Chemokine Signaling Pathway, Antigen Processing and Presentation, and Fc epsilon RI Signaling Pathway further supported the role of an abnormal immune microenvironment, potentially hindering the successful implantation [45,46]. Also there were significantly down regulated pathways which included Terpenoid Backbone Biosynthesis, Biosynthesis of Unsaturated Fatty Acids, Butanoate and Propanoate Metabolism, Mucin Type O-Glycan Biosynthesis, and Steroid Biosynthesis. These pathways had genes namely ACAT1, ACAT2, HADHA, ECHS1, GALNT family members, and CYP family genes which were commonly associated across these pathways focusing on suppression of lipid and energy metabolism, glycan modification, and steroid hormone synthesis, all of which are critical for maintaining a receptive endometrial environment [47–49]. In a reproductive context, terpenoid pathway could affect the local synthesis of steroids or important implantation factors (since cholesterol is a precursor for estradiol and progesterone synthesis in peripheral tissues, and is also crucial for cell membrane fluidity and signaling molecule rafts [50]. Butanoate metabolism, involving short-chain fatty acid utilization, suggests alterations in how cells handle certain substrates and could be linked to the energy metabolism of the tissue or even the influence of microbial metabolites. Though butyrate is classically a gut microbial product, human endometrial cells do express enzymes for short-chain fatty acid metabolism; a suppression of this pathway might reflect lower oxidative metabolic activity (since butyrate metabolism feeds into acetyl-CoA and the TCA cycle) [51].

#### 3.4.3 Gene Ontology (GO) Analysis

Gene ontology enrichment analysis categorized gene functions into three major categories - Biological Processes (BP), Molecular Functions (MF), and Cellular Components (CC) **Figure 4**. For the biological processes the genes were enriched in metabolic regulation and biosynthetic processes, including regulation of metabolic process, regulation of macromolecule biosynthetic process, regulation of cellular metabolic process. **Figure 4 B**. For cellular components the genes were enriched in Vesicle lumen, cytoplasmic vesicle lumen, and secretory granule lumen, “early endosome and blood microparticle” **Figure 4 C**. And for molecular functions the genes were enriched in “Potassium ion transport activity”, “monocarboxylic acid binding activity”, “Transmembrane transporter activity” and, “Retinoic acid and retinoid binding” **Figure 4 D**.

**Figure 4A.**
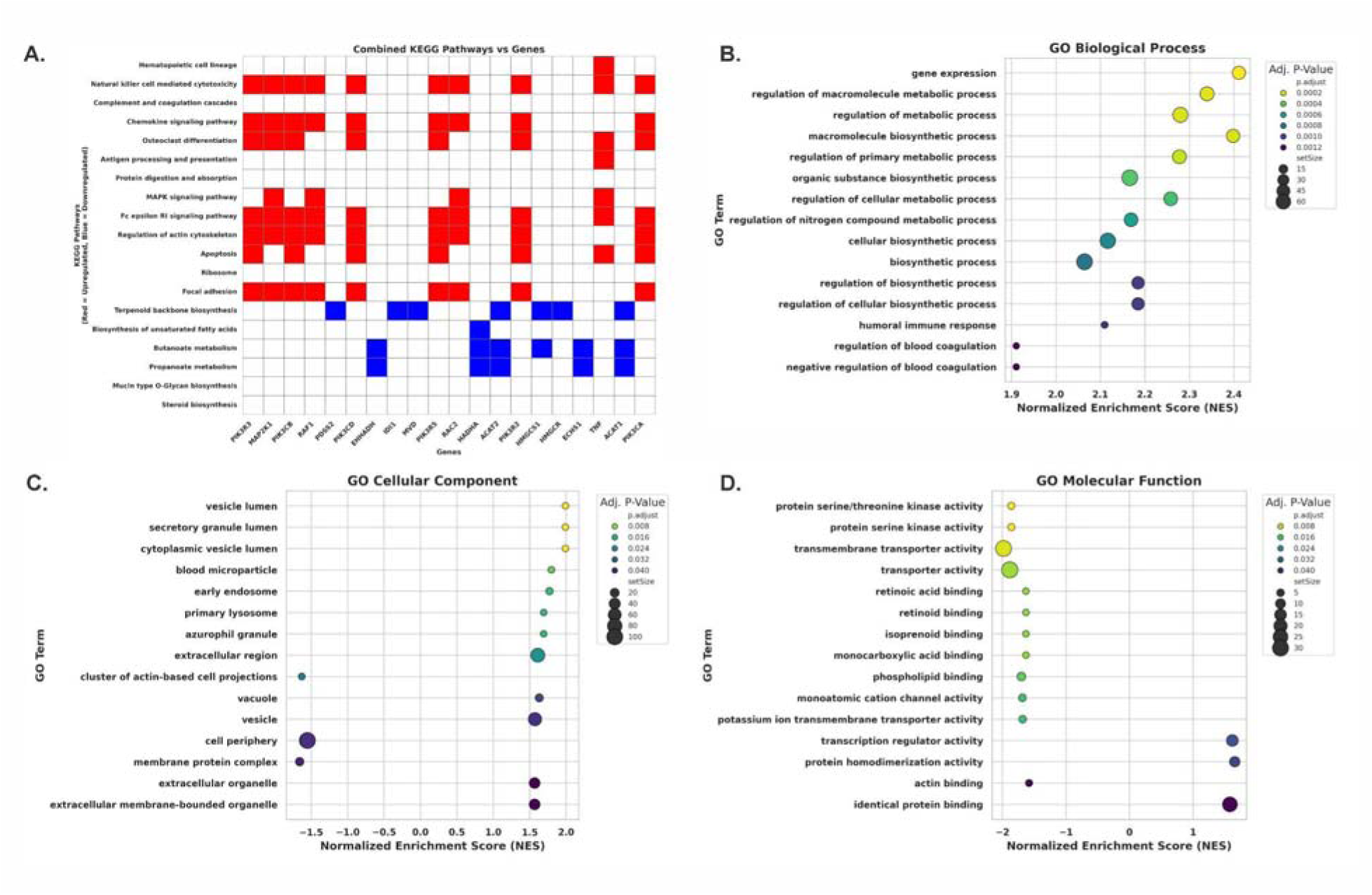
Heatmap representing DEGs in KEGG pathways. Red colour represents upregulated genes and blue represents downregulated genes **B.** GO Biological Process enrichment dot plot showing significantly enriched pathways (y-axis) ranked by normalized enrichment score (x-axis). Dot size indicates gene set size, and color represents adjusted p-value. **C.** GO Cellular Component enrichment results showing subcellular localization of DEGs. **D.** GO Molecular Function enrichment dot plot highlighting activities such as serine/threonine kinase activity, transmembrane transporter activity, and binding functions (e.g., retinoid, actin). NES and p-values guide significance.

**Figure 5A.**
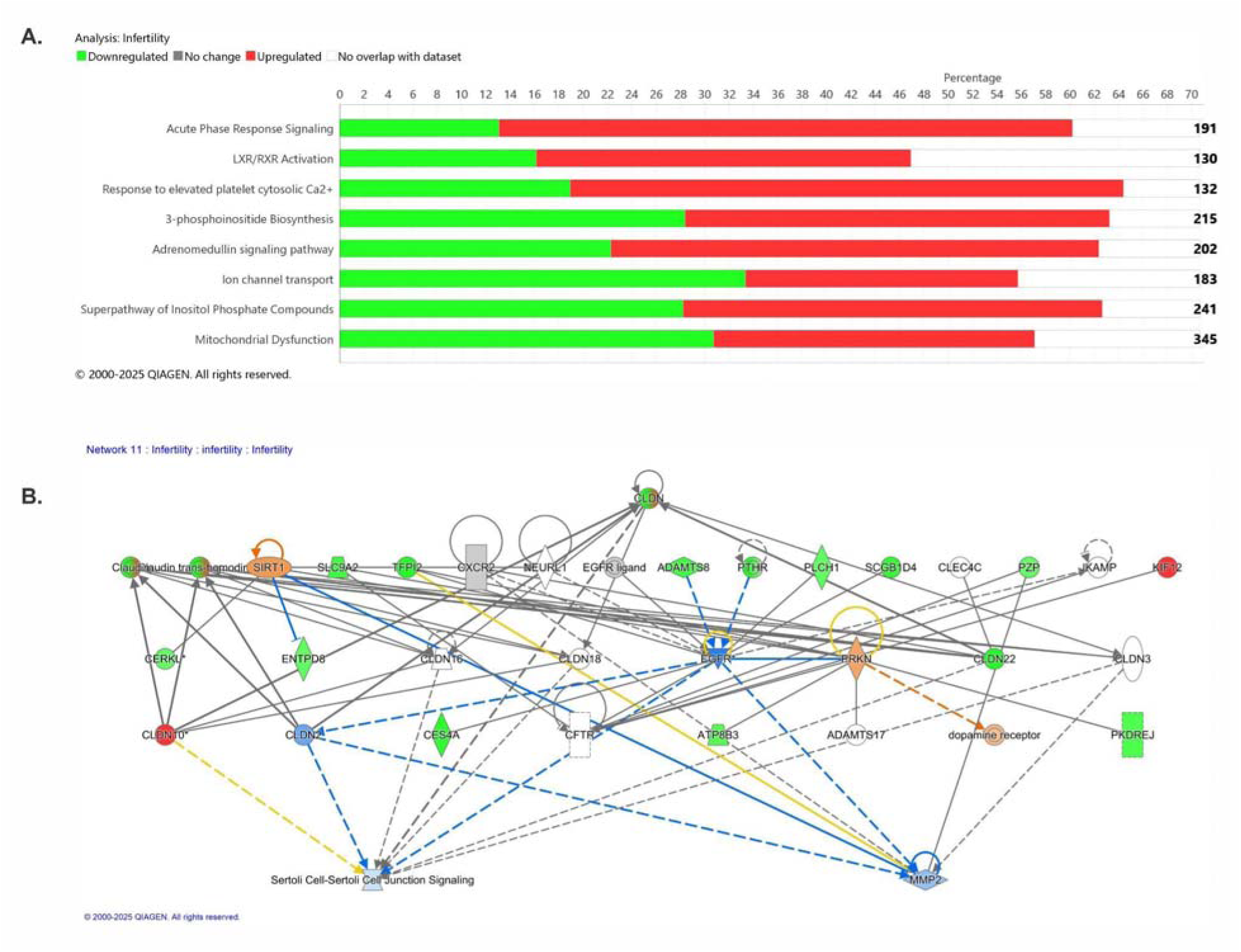
Bar plot representing top canonical pathways significantly impacted in infertility, as identified by IPA. The colour highlights the proportion of genes in each pathway that are upregulated (red), downregulated (green), or unchanged (grey) is depicted. **B.** Network showing the functional connectivity of genes involved in “Sertoli Cell-Sertoli Cell Junction Signaling.” The colour of the nodes represents change in gene expression, red for upregulated, green for downregulated, and white for unchanged. Edges indicate known interactions (solid lines for direct, dashed lines for indirect), with blue and orange lines representing predicted inhibition or activation, respectively.

### 3.5 Ingenuity Pathway Analysis (IPA)

#### 3.5.1 Canonical Pathways

The canonical pathways from the IPA analysis included both activated and inhibited pathways. The activated pathways included “Acute Phase Response Signaling” which involved genes such as *C3, C4BPA, FGA, FGB, IL6ST, ORM1, ORM2,* and *SOD2*. This supports the role of systemic inflammation and innate immune modulation during implantation [52]. LXR/RXR Activation, was another activated pathway. It involved genes such as ORM1, ORM2, PTGS2, and FGA known as lipid regulators, which plays an important role in cholesterol homeostasis and membrane remodeling during endometrial receptivity [52]. It was also found that “Neutrophil Degranulation”, involving genes such as ATP8A1, C3, CAMP, CD68, CDA, ORM1, ORM2, SLC15A4, was also activated, stating the significance of neutrophil-mediated inflammation in the endometrial environment. Response to Elevated Platelet Cytosolic Ca² involving, genes FGA, FGB, ORM1, ORM2 was also activated, pointing towards dysregulated in platelet function relevant to implantation-related vascular modulation [53].

There were also a number of metabolic and regulatory signaling pathways that were inhibited. Inhibited pathways included “Superpathway of Inositol Phosphate Compounds, and 3-Phosphoinositide Biosynthesis”. These involved genes like ACP6, ALPL, PIK3C2G, PIP5K1B, PLCH1, PPP1R14C, PSPH, which play central roles in calcium signaling and membrane integrity [54]. The “Adrenomedullin Signaling Pathway” was also inhibited, involving genes namely C3, GAD1, GPR37, PIK3C2G, PLCH1, highlighting dysregulation in angiogenesis and endometrial vascularization [55]. Ion Channel Transport was suppressed, with genes ATP12A, ATP6V0E2, ATP8A1, ATP8B3, CAMK2B, TRPM6, TTYH2, WNK4 emphasising on disturbances in epithelial polarization and ion balance, essential for embryo adhesion [56]. Most critically, “Mitochondrial Dysfunction” was inhibited. This pathway involved genes namely ATP12A, CAMK2B, CAPN6, CREB3L1, GPX3, HAP1, PIK3C2G,

PPARGC1A, SOD2, pointing towards impaired oxidative stress regulation and energy metabolism, all of which are important factors for implantation [57]. Key regulators of mitochondrial biogenesis such as PGC-1α (PPARGC1A) were downregulated in our dataset, consistent with a state of reduced mitochondrial function. In case of infertility, an elevated acute phase response means that the endometrium is responding as if under attack or stress, which could be due to low level inflammation impaired [58]. Also, the activation of LXR/RXR (liver X receptor/retinoid X receptor) pathways points to altered lipid metabolism and inflammatory regulation. LXR/RXR are nuclear receptors that *“regulate lipid synthesis and transport, induce genes involved in cholesterol transport, and modulate immune and inflammatory responses”*. Their activation may indicate a compensatory response to inflammation. For example, LXR activation can inhibit inflammatory gene expression in macrophages and promote cholesterol efflux. This could be a way to reduce the inflammation or deal with lipid by-products of tissue stress. Thus, the upregulation of LXR/RXR and acute phase pathways together paint a picture of an endometrium in an “alert” state: metabolically reprogrammed and mounting an innate immune response [59].

#### 3.5.2 Upstream Regulators

IPA identified several upstream regulators which were then cross-referred with the Comparative Toxicogenomics Database (CTD); all identified upstream regulators showed only hypothetical associations with infertility. None had known therapeutic targets (T) or markers/mechanisms (M) for the disease. Amongst key activated regulators were PRKAA1 (AMP-activated protein kinase α1). It modulated key metabolic and inflammatory genes such as PPARGC1A, PLA2G2A, and SOD2, which are recurrently involved in KEGG pathways (e.g., obesity, glucose metabolism), canonical pathways (e.g., mitochondrial dysfunction), and diseases like carbohydrate imbalance and gut inflammation. AMPK is activated under conditions of low energy (high AMP/ATP ratio) and functions to restore energy balance by initiating catabolic processes and inhibiting heavily energy-dependent anabolic pathways. This analysis suggests that AMPK is significantly engaged in the endometrium of unexplained infertility patients through *PRKAA1* **[60]**. Activation of HDAC4, a transcriptional repressor involved in chromatin remodeling, affected the expression of IGFBP1, NR4A2, PTGS2, and GADD45A genes linked to apoptosis, oxidative stress, and embryonic development, often mapped across conventional and KEGG pathways. Another active regulator, PTEN, targeted metabolic and immunological genes (C3, GPX3, CXCL14, PPARGC1A, PTGS2), suggesting endometrial receptivity-related immunometabolic homeostasis. Although acknowledged for DNA repair, BRCA1 was activated linked to oxidative stress and developmental regulators such GADD45A, CAMK2B, HSD11B2, and NEDD9. These regulators target DEGs such PTGS2, PPARGC1A, SOD2, CXCL14, and NR4A2, which have roles in mitochondrial malfunction, apoptosis, neutrophil degranulation, and glucose metabolism. PTEN (Phosphatase and Tensin Homolog) is a tumor suppressor and a strong blocker of the PI3K/AKT signaling pathway. In the physiology of the endometrium, PI3K/AKT signaling is very important for cell survival, growth, and decidualization, which is when stromal cells change to get ready for implantation. For an endometrium to be receptive, it needs to have a balanced decidualization response. Problems with decidualization have been linked to implantation failure, repeated pregnancy loss, and unexplained infertility **[61]**. **Suppressed** Regulators Indicate Protective Mechanism Loss: The transmembrane anti-inflammatory receptor IL10RA and downstream genes C3, CA2, PPARGC1A, and SORBS2 were dramatically suppressed. Pro-inflammatory mechanisms like acute phase response and GI inflammation are supported by this. Most commonly IL-10 is known as a tolerogenic or immunosuppressive cytokine as it plays a critical role in pregnancy for keeping in check of harmful immune responses **[62]**. Elevated IL-10 is associated with successful pregnancy, whereas deficiencies in IL-10 signaling are linked to pregnancy complications and immune-mediated implantation failures. The identification of IL10RA as one of the significant regulators suggests that there is deficit in the IL-10 signaling axis in the infertile endometrium and its inhibition justifies that there are gene expression changes consistent with a state of reduced IL-10 activity.

RAD51, a critical DNA repair player, was also decreased, which may compromise genomic stability in endometrial cells under oxidative stress, supporting mitochondrial dysfunction. MYC, FOXA1, ERG, and EGFR regulated cell proliferation, differentiation, and survival, affecting gene sets involved in development, angiogenesis, and immunological homeostasis like LAMB3, GADD45A, PTGS2, HSD11B2, and NTRK2. Inhibition of RAF1 and CASR also reduced MAPK signaling, vascular signaling, and inflammatory resolution, consistent with downregulated canonical and KEGG pathways such adrenomedullin signaling and ion channel trafficking. PTGS2 (COX-2) was identified as a highly associated gene which was targeted by 9 different regulators, and involved across inflammatory, metabolic, and vascular pathways. PPARGC1A and SOD2, both crucial for mitochondrial and glucose metabolism, were modulated by multiple regulators (PTEN, PRKAA1, MYC, IL10RA), strengthening their role in the infertile endometrial phenotype. Although none of these upstream regulators are currently listed as direct marker or therapeutic genes in the CTD for female infertility, their modulation of key DEGs repeatedly observed across major pathway clusters points to their emerging relevance as upstream control nodes in the pathogenesis of unexplained infertility.

#### 3.5.3 Diseases or BioFunctions Annotation

The disease and function annotation analysis identified 9 activated and 4 inhibited biological pathways, many of which are not discussed in the context of infertility. Among the most significantly enriched annotations were “Inflammation of the gastrointestinal tract”, “Gastroenteritis”, “Abnormality of large intestine”, “Enteritis”, “Colitis”. These pathways shared a robust set of 25-33 genes (e.g., AOX1, ATP12A, C3, FGA, PTGS2, IL6ST, SLC15A4 etc) known to be involved in immune signaling, epithelial stress, and metabolic reprogramming. While these GI-related diseases are not typically linked to female infertility, emerging research suggests that systemic or local gut inflammation may disrupt reproductive homeostasis through the gut-endometrium axis, affecting cytokine levels, endometrial receptivity, and immune tolerance during implantation.

Transport of ion, Transport of monovalent inorganic cations were amongst downregulated pathways, involving genes such as ATP8A1, CAMK2B, SLC15A2, TRPM6, WNK4, essential for fluid homeostasis and epithelial polarization, which plays a crucial for successful implantation. Additionally, the suppression in function cell movement of sperm was observed which supported the dysregulation of suppression of ion transport and mitochondrial pathways observed earlier, highlighting the possibility of affecting fertility outcome through impaired sperm-endometrium signaling. Carbohydrate metabolism-related annotations were also activated which included Quantity of carbohydrate, Concentration of D-glucose, and Obesity. These reflect a shift toward metabolic stress and insulin resistance, hallmarks of PCOS and obesity-associated infertility. Genes such as PPARGC1A, FMO5, SOD2, CXCL14, CYP26A1, PTGS2 were frequently repeated in these pathways and are known regulators of glucose handling and oxidative stress in reproductive tissues.

#### 3.5.4 Novel Functional Associations and Network Insights

Disease and biofunction annotations highlighted two novel statistical and biologically significant insights, which are GI inflammation and large intestine abnormalities. These diseases are well-studied in gastroenterology, but their link to female infertility is unknown. The genes (C3, PTGS2, IL6ST, ATP12A, NNMT) associated with these annotations support the canonical pathway analysis of inflammatory and metabolic signatures. This overlapping highlights that persistent GI inflammation may modify systemic immune and metabolic conditions, reducing endometrial receptivity and implantation success. Further, analysis of gene interaction networks for Endocrine System Disorders, Gastrointestinal Disease, and Organismal Injury and Abnormalities was done. Despite considerable dysregulation in our dataset, Two gene entries: CLDN (claudin) and PTHR (parathyroid hormone receptor) in the network did not have therapeutic or mechanistic roles in the CTD, suggesting their potential uniqueness in infertility. After reviewing literature, it was found that CLDN1 and CLDN4, members of the claudin family, have been linked to embryo implantation and endometrial barrier modulation, but the generic CLDN group, as annotated in IPA, has never been linked to infertility. This suggests that the present network’s claudin isoforms (e.g., CLDN2, CLDN10, CLDN22, CLDN18) may participate in novel tight junction signaling events related to reproductive failure. Although engaged in calcium signaling and hormonal pathways, PTHR has no direct or inferred connections with infertility in existing literature or databases, suggesting an uncharacterized role that needs mechanistic validation. The network analysis (Figure 6) shows EGFR and MMP2 as central nodes connecting CLDN isoforms and associated regulators like PRKN, SIRT1, and TFPI2, suggesting that infertile endometrium may disrupt tight junction remodeling, epithelial integrity, and proteolytic balance. CES4A, SLC9A2, and SCGB1D4 were also considerably downregulated and connected to epithelial barrier function, transporter activity, and immunological modulation, suggesting tissue homeostasis breakdown.

**Figure 6:**
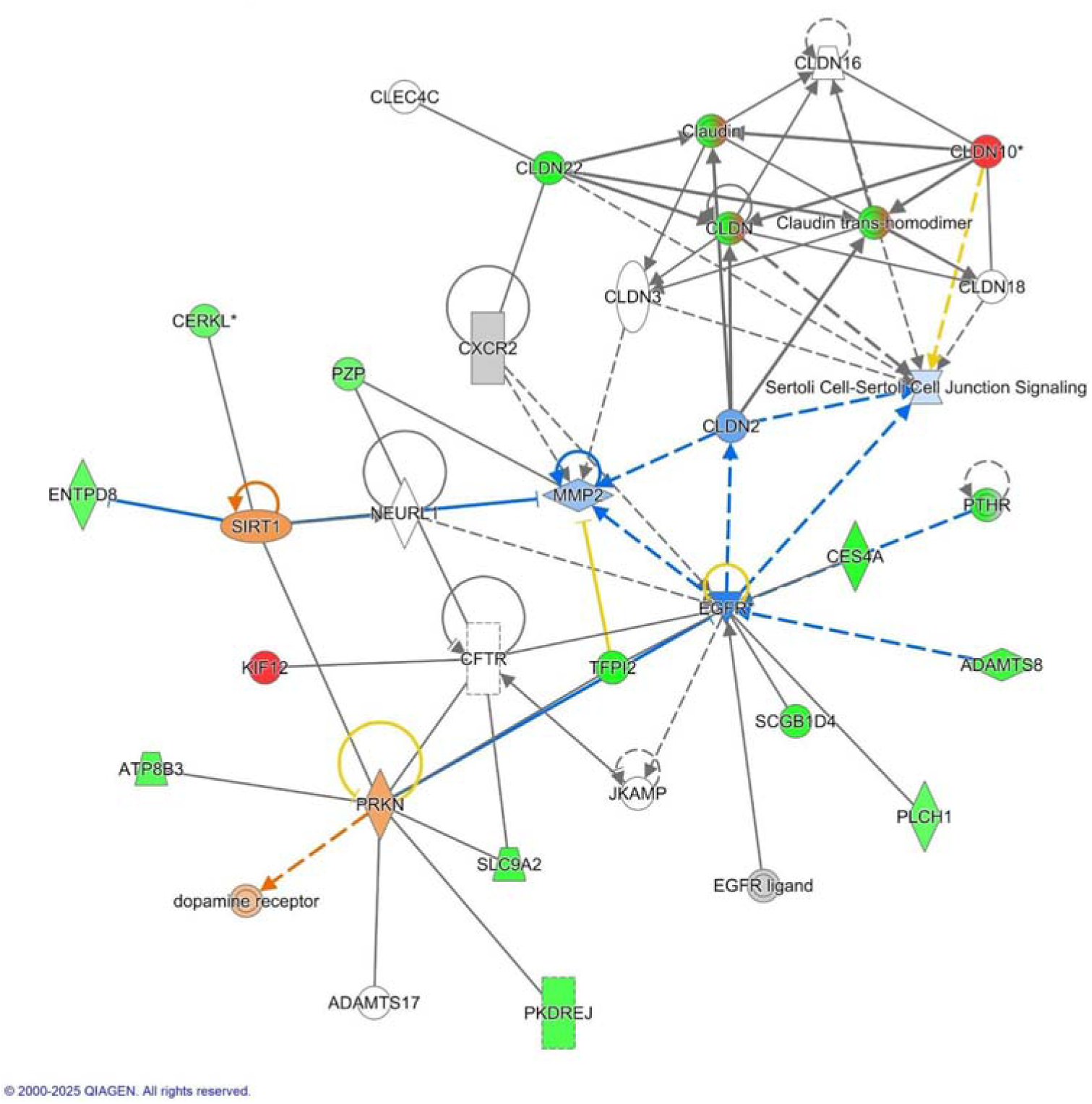
IPA network showing isoforms of CLDN responsible for novel tight junction signaling events related to reproductive failure.

#### 3.5.5 Protein–Protein Interaction (PPI) Network

To better understand the functional relationships among the differentially expressed genes (DEGs), a Protein–Protein Interaction (PPI) network in figure 4.6 was constructed using the STRING database. The network consisted of 164 nodes and 18 edges, indicating significant enrichment (p = 9.11 × 10 ). Protein–protein interaction network analysis of the novel infertility-associated genes revealed a highly structured and biologically coherent network rather than random gene associations, supporting the functional relevance of these previously unreported candidates. The network was dominated by genes involved in epithelial ion transport, solute exchange, extracellular nucleotide metabolism, and GPCR-mediated signaling, highlighting epithelial homeostasis and immunometabolic regulation as central features of infertility-associated endometrial dysfunction.

A prominent network module comprised multiple solute carrier (SLC) transporters (SLC1A1, SLC7A1, SLC15A2, SLC43A1, SLC9A2) together with ion regulators such as WNK4, TRPM6, ATP12A, ATP6V0E2, and KCNJ2. These genes collectively regulate sodium, potassium, proton, magnesium, and amino acid transport, which are critical for maintaining endometrial epithelial polarity, luminal fluid composition, and pH balance. Proper ionic and nutrient gradients are essential for embryo apposition and implantation; therefore, dysregulation of this transport network suggests a uterine environment that is physiologically imbalanced and hostile to embryo attachment.

A second highly interconnected cluster centered on purine and nucleotide metabolism genes, including ENPP3, ENTPD8, UCK2, GMPR, and CDA. These genes regulate extracellular ATP degradation and intracellular nucleotide recycling, processes that are tightly linked to cellular stress responses and immune signaling. Extracellular ATP acts as a danger-associated molecular pattern (DAMP) that activates innate immune responses; thus, altered purinergic signaling may amplify local inflammation and disrupt immune tolerance at the maternal–fetal interface. The convergence of metabolic and immune signaling within this module reinforces the concept of infertility as an immunometabolic disorder rather than a purely hormonal condition.

GPCR and neuromodulatory signaling pathways were also represented within the network, with genes such as GPR37, GPR18, ADGRF1, OPRPN, and PTH2R forming a distinct signaling axis. These receptors are known to mediate neuroimmune communication, stress sensing, and hormonal responsiveness. The presence of neuronal differentiation–associated genes such as NEUROG2 and calcium-binding proteins such as HPCAL4 further supports the emerging role of neuroendocrine-like signaling in endometrial function. This finding aligns with growing evidence that the endometrium integrates neuronal, immune, and metabolic cues to regulate receptivity and implantation timing.

Additional network components included epithelial differentiation and barrier-related genes such as KRT8, LAMB3, CLDN22, SERPINB5, UPK1B, and PROM2, indicating altered epithelial integrity and extracellular matrix interactions. Disruption of epithelial cohesion and barrier function is known to impair embryo adhesion and invasion, suggesting that structural remodeling contributes to the non-receptive endometrial phenotype observed in infertility.

Importantly, immune-modulatory genes such as VTCN1, KIR3DL3, ACKR1, and BANK1 were also integrated into the network, indicating dysregulated immune tolerance mechanisms. VTCN1 (B7-H4) functions as an immune checkpoint molecule, while KIR3DL3 is involved in NK cell regulation, supporting the presence of an immune environment that is active yet improperly regulated. This finding is consistent with the observed upregulation of innate immune pathways and loss of anti-inflammatory signaling in infertile endometrium.

## Discussion

This study integrates differential gene expression analysis, gene ontology and KEGG enrichment, Ingenuity Pathway Analysis (IPA) and weighted gene co expression network analysis (WGCNA) to show that unexplained female infertility is characterised by several changes in the endometrium.

First, we observed **elevated innate immune and inflammatory activity** through pathways such as natural killer (NK) cell mediated cytotoxicity, complement and acute phase signalling, and neutrophil degranulation. Second, there was a marked **suppression of metabolic capacity**, including lipid and steroid biosynthesis, short chain fatty acid metabolism and mitochondrial functions. Third, we found an unreported **link to gastrointestinal inflammatory signatures**, suggesting that gut derived immune or metabolic signals may influence endometrial receptivity.

The Network-level findings from WGCNA highlighted a turquoise module whose hub genes include *PTGS2*, *SOD2* and *PPARGC1A*. All of these genes are responsible in impaired endometrial receptivity. The emergence of *C7orf50* (cholesin), a cholesterol related factor, as a negatively associated gene further associates metabolic status of the system to implantation biology.

### Immune activation and inflammatory signalling

The findings from our enrichment analyses point to a pro inflammatory state in the infertile endometrium. Amongst the up regulated pathways were natural killer (NK) cell mediated cytotoxicity [63] and complement/Coagulation Cascades [46]. While uterine NK cells normally assist implantation, the presence of excessive NK cell activity or unchecked complement activation could damage the embryo or disrupt maternal fetal tolerance. Up regulation of acute phase response genes and neutrophil degranulation supports the idea that the endometrial tissue is reacting as if it were injured or infected. Consistent with this, we noted down regulation of *IL10RA*, the receptor for the anti inflammatory cytokine interleukin 10; loss of IL 10 signalling may tilt the balance toward inflammation and affect receptivity.

### Metabolic and energy deficits

In our analysis we also observed suppression of metabolic and energy producing pathways. The down regulated genes were involved in cholesterol and steroid synthesis, unsaturated fatty acid production, and hort chain fatty acid metabolism. Mitochondrial dysfunction emerged as a top inhibited pathway, stating that energy production and reactive oxygen species handling are compromised. *PPARGC1A*, the gene responsible for regulation of mitochondrial biogenesis, was reduced, and antioxidant genes such as *SOD2* and *GPX3* were up regulated, indicating a compensatory response to oxidative stress. The terpenoid backbone pathway, responsible for synthesis of cholesterol for estradiol and progesterone synthesis, was also suppressed. Reduced cholesterol synthesis could limit local hormone signalling and alter membrane fluidity, making it harder for the embryo to attach.

### Upstream regulators and DNA damage

We identified several upstream regulators from Ingenuity Pathway Analysis. Activated *PRKAA1* (AMP activated protein kinase α1) suggests the endometrium senses low energy and shifts toward catabolic processes. Down reguled *IL10RA* confirms impaired anti inflammatory control, while reduced *RAD51* points to an inability for DNA repair. Persistent inflammation and oxidative stress can lead to DNA damage and cellular senescence, which have been implicated in implantation failure. The interconnection of these regulators states a situation of an endometrium under metabolic and inflammatory stress. Amongst all, RAD51 (RAD51 Recombinase) is an enzyme that plays a critical role in the homologous recombination repair of DNA double-strand breaks. Its role in animals has been established but not in human infertility. Its presence as an upstream regulator indicates that DNA damage responses could be altered in these endometrial samples. The two well known inducers of DNA damage are Chronic inflammation and oxidative stress. The inhibition of RAD51 suggests an accumulation of DNA damage or a stress response where the cell is not able to regulate repair pathways, associating it with cellular senescence. Cells under the state of senescence accumulate in aging and in inflammatory conditions and secrete pro-inflammatory cytokines that can worsen tissue inflammation.

### A possible gut–endometrium axis

The output of IPA disease and function marked a connecting link with inflammatory disorders outside the reproductive system that was one of the most interesting insights. It focused on gastrointestinal (GI) inflammation and highlighted diseases like inflammation of gastrointestinal tract and abnormality of the large intestine as important links. Recent reviews on the gut–reproductive axis explain that a healthy gut microbiota is closely linked to immune homeostasis, whereas gut dysbiosis can trigger immune dysfunction and chronic low grade inflammation. Such inflammation is associated with disorders like endometriosis, polycystic ovary syndrome, insulin resistance and obesity, all the conditions that often accompany infertility. Reduced microbial diversity in both the gut and reproductive tract can impair immunosurveillance and disrupt hormone signalling. Moreover, dysbiosis may limit nutrient absorption, thereby affecting the supply of vitamins and macronutrients needed for hormone production and energy metabolism [64]. Chronic low grade inflammation can alter ovarian function and oocyte quality, and leaky gut associated translocation of bacterial products into the bloodstream may activate uterine immune cells, further compromising receptivity [65]. Our identification of *C7orf50* (cholesin), a gut derived hormone that suppresses the hepatic cholesterol regulator SREBP 2 [66], strengthens this gut–endometrium connection; altered intestinal cholesterol signalling could influence local steroid hormone availability in the endometrium.

### Novel genes and epithelial barrier functions

In our network analysis we also uncovered new genes involved in epithelial barrier integrity. Claudin family members (CLDN genes) form tight junctions that maintain epithelial polarity were highlighted in network analysis of IPA. In the uterine epithelium, claudin 4 expression normally increases during implantation, and women with unexplained infertility show higher claudin 4 levels compared with fertile individuals. Also the reduced claudin 3 and claudin 4 expression is associated with endometriosis. The down regulation of ion transporters such as *WNK4* and *TRPM6* in our analysis suggests disturbances in epithelial polarization and fluid balance. Together, these observations point out that tight junction remodelling and impaired ion transport may compromise the luminal microenvironment required for embryo implantation. *PTHR* (parathyroid hormone receptor) also emerged in our network, yet little is known about its role in fertility; further work is needed to determine whether it participates in calcium dependent signalling during implantation.

### Epithelial-immune–metabolic dysregulation

The interaction network reveals that the novel infertility-associated genes converge on an interconnected biological framework involving epithelial ion homeostasis, purinergic signaling, neuroimmune communication, and immune tolerance. Rather than representing isolated molecular abnormalities, these genes form an integrated stress-response network that reflects chronic immunometabolic imbalance, epithelial dysfunction, and disrupted signaling at the maternal–fetal interface.

### Clinical implications and future directions

These findings combined portray a scenario that there are several factors behind unexplained female infertility. Ihe factors may include a combination of excessive innate immune activation, suppressed metabolic activity and a systemic inflammatory state possibly rooted in the gut. Addressing these issues may therefore require a holistic approach. Strategies to minimize local inflammation and restore IL 10 signaling, and to promote metabolic health, such as improving diet, managing insulin resistance, and boosting mitochondrial function, could make the endometrium more receptive. Because of the new connection between the gut and reproduction, it is worth looking into treatments that target gut dysbiosis, such as probiotics, dietary fiber, and anti-inflammatory diets. . Finally, the novel genes identified here (including *C7orf50* and specific claudin isoforms) provide potential biomarkers for diagnosing endometrial dysfunction and may serve as entry points for new treatments.

## Conclusion

In conclusion, the convergence of evidence in our study - from canonical pathways and KEGG, to upstream regulators and disease associations - all supports a model of unexplained infertility rooted in an imbalanced endometrial environment. There is excessive inflammation and immune activation (likely contributing to implantation failure by attacking or not properly nurturing the embryo), coupled with oxidative stress and suppressed cellular metabolism (leading to an endometrium that is underprepared energetically and structurally for pregnancy). These findings not only enhance our understanding of the biological underpinnings of unexplained infertility but also open new directions for both research and clinical management. Targeting the highlighted pathways (inflammation, oxidative stress, metabolic support) and exploring the systemic connections (such as the gut-uterus axis) represent promising frontiers to transform “unexplained” infertility into a tractable condition with clear diagnostic markers and tailored treatments. Ultimately, by addressing the root causes identified, calming the inflammatory storm and reawakening the metabolic vigor of the endometrium may improve the chances for successful implantation and healthy pregnancies in this perplexing group of patients.

## Declarations

## Supporting information

https://docs.google.com/document/d/1GI6tYFj_wQyU3uO2r44K2myeRFkQ8hdQ/edit?usp=drive_link&ouid=100845841708565570348&rtpof=true&sd=true

## Acknowledgements

The authors would like to thank the Centre for Systems Biology and Bioinformatics and the Department of Biotechnology, Panjab University, Chandigarh, for providing the computational and infrastructural support required to carry out this study.

## Author Contributions

**Ritika Patial:** Conceptualization, Data curation, Formal analysis, Methodology, Investigation, Writing Original Draft. **Sonalika Ray:** Data analysis, Results compilation and interpretation, Review & Editing

**Dr. Kashmir Singh:** Project administration, Supervision, Methodological guidance, Review.

**Prof. R.C. Sobti**: Project administration, Supervision, Review & Editing.

## Data Availability

The gene expression dataset analyzed in the study is publicly available in the NCBI Gene Expression Omnibus (GEO) repository under accession number GSE92324.

## Ethics Approval and Consent to Participate

This study is based on the re-analysis of publicly available transcriptomic data and does not involve any new human or animal subjects. Therefore, ethics approval and participant consent were not required.

## Supplementary

We found 3 activated and 5 significantly inhibited canonical pathways Table (1).

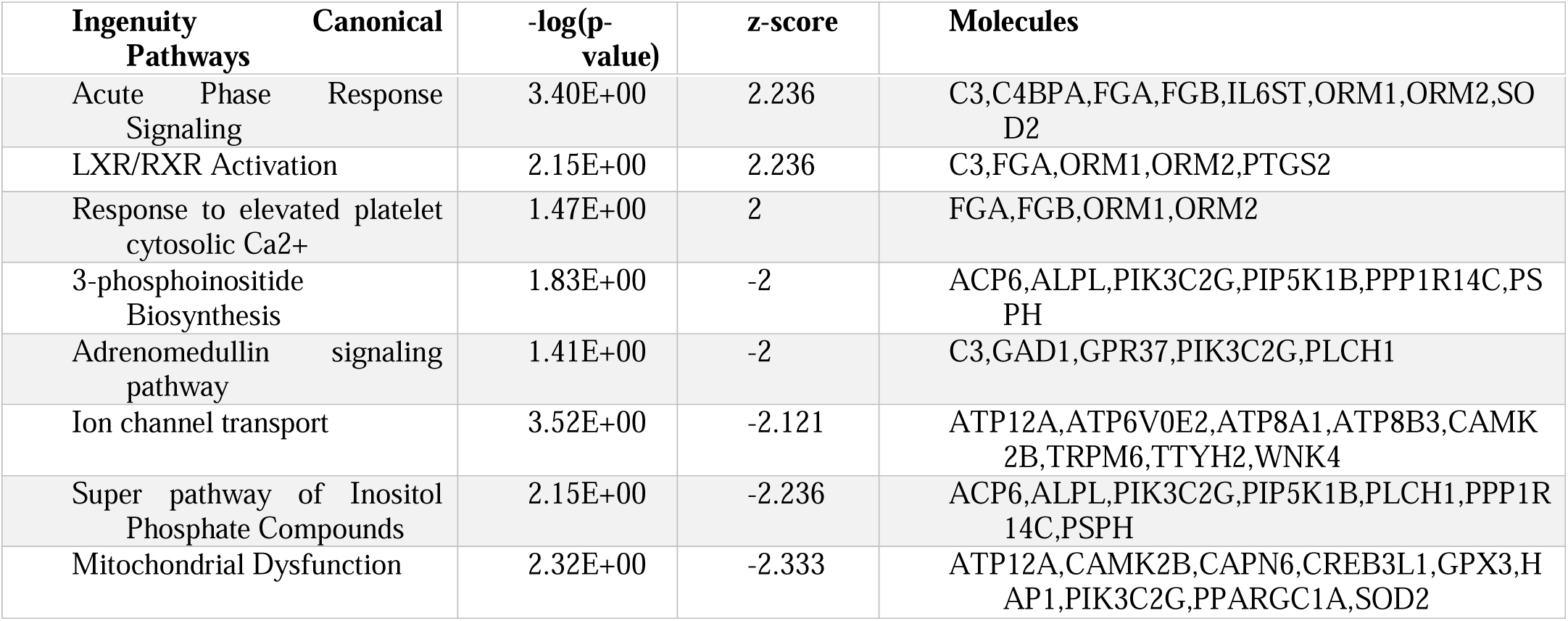

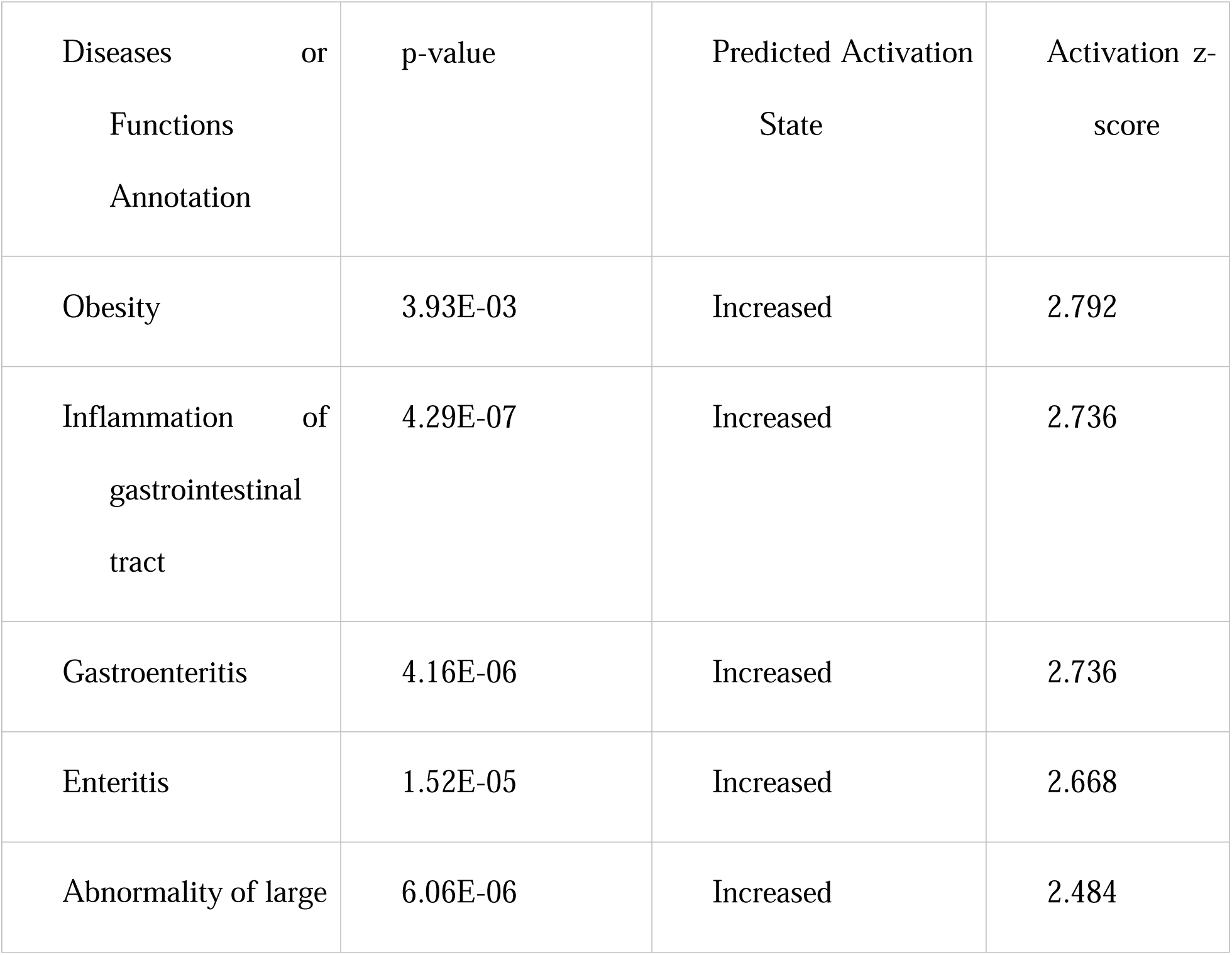
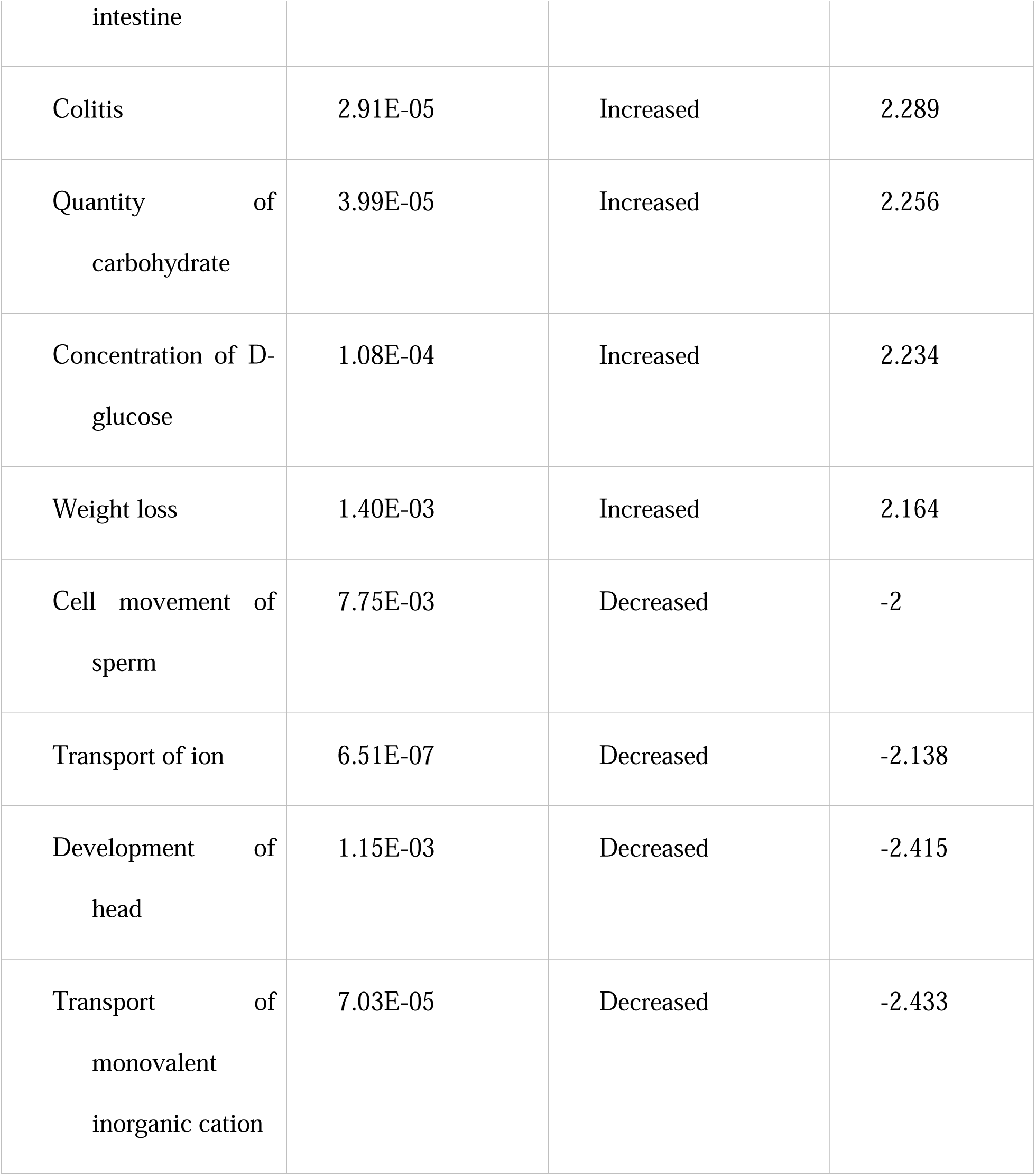

## References

[1] H. Zhu, X. Zhou, R. Li, Q. Gao, X. Wang, P. Cheng, R. Liu, C. Yin, Y. Hao, Global prevalence of infertility: a systematic review and meta-analysis of Community-based studies, (2023). 10.22541/au.169535894.43892783/v1.

[2] Y. Liang, J. Huang, Q. Zhao, H. Mo, Z. Su, S. Feng, S. Li, X. Ruan, Global, regional, and national prevalence and trends of infertility among individuals of reproductive age (15–49 years) from 1990 to 2021, with projections to 2040, Hum. Reprod. 40 (2025) 529–544. 10.1093/humrep/deae292.

[3] S. Ara Anwary, M.N.I. Mondal, M.R. Karim, M.M. Rahman, M. Alfazzaman, Z. Mahzabin, M.M. Rahman, A. Nahar, Causes of Infertility among the Couples Who are Attending the Infertility OPD of BSMMU, Med. Today 35 (2023) 20–26. 10.3329/medtoday.v35i1.64934.

[4] R. Heidarzadehpilehrood, M. Pirhoushiaran, M. Binti Osman, H. Abdul Hamid, K.-H. Ling, Weighted Gene Co-Expression Network Analysis (WGCNA) Discovered Novel Long Non-Coding RNAs for Polycystic Ovary Syndrome, Biomedicines 11 (2023) 518. 10.3390/biomedicines11020518.

[5] Phillips, K., Olanrewaju, R. A., & Omole, F. (2023). Infertility: Evaluation and Management. American Family Physician, 107(6), 623–630., (n.d.).

[6] N. Joshi, J. Chan, Female Genomics: Infertility and Overall Health, Semin. Reprod. Med. 35 (2017) 217–224. 10.1055/s-0037-1603095.

[7] Wikipedia contributors. (2024, April 10). Fragile X-associated primary ovarian insufficiency. In Wikipedia. https://en.wikipedia.org/wiki/Fragile_X-associated_primary_ovarian_insufficiency, (n.d.).

[8] S. Siddharth, B. Farooq, N. Kumar, M. Burhan, Effect of Lifestyle in Female Infertility: A Review Based Study, Int. J. Res. Appl. Sci. Eng. Technol. 11 (2023) 1777–1783. 10.22214/ijraset.2023.56307.

[9] J. Li, Y. Huang, S. Xu, Y. Wang, Sleep disturbances and female infertility: a systematic review, BMC Womens Health 24 (2024) 643. 10.1186/s12905-024-03508-y.

[10] Zhu, M., Yi, S., Huang, X., Meng, J., Sun, H., & Zhou, J. (2020). Human chorionic gonadotropin improves endometrial receptivity by increasing the expression of homeobox A10. Molecular human reproduction, 26(6), 413–424. 10.1093/molehr/gaaa026, (n.d.).

[11] F. Dong, W. Zhang, B. Sun, W. Niu, J. Zhai, Y. Guo, F. Wang, Integrated Transcriptome and Proteome Analyses Reveal Differentially Expressed Genes and Proteins in Granulosa Cells from Female Patients with Metabolic Syndrome-associated Infertility, Curr. Med. Chem. 32 (2025). 10.2174/0109298673357582241223070335.

[12] C.M. Da Luz, M.G. Da Broi, L.D.O. Koopman, J.R. Plaça, W.A. Da Silva-Jr, R.A. Ferriani, J. Meola, P.A. Navarro, Transcriptomic analysis of cumulus cells shows altered pathways in patients with minimal and mild endometriosis, Sci. Rep. 12 (2022) 5775. 10.1038/s41598-022-09386-4.

[13] E. Vargas, A. Sola-Leyva, N.M. Molina, S. Bonilla, E. Salas-Espejo, S. Ruiz-Durán, L. Terrón-Camero, I. Leonés-Baños, L. Martinez-Navarro, R. Sánchez, B. Romero-Guadix, J. Mozas, L. Aghajanova, E. Andrés-León, S. Altmäe, P-366 Unraveling the whole transcriptome profiles of receptive phase endometrium in infertile women, Hum. Reprod. 38 (2023) dead093.724. 10.1093/humrep/dead093.724.

[14] B.N. Bui, V. Kukushkina, A. Meltsov, C. Olsen, N. Van Hoogenhuijze, S. Altmäe, F. Mol, G. Teklenburg, J. De Bruin, D. Besselink, L. Stevens Brentjens, D. Obukhova, M. Zamani Esteki, R. Van Golde, A. Romano, T. Laisk, G. Steba, S. Mackens, A. Salumets, F. Broekmans, The endometrial transcriptome of infertile women with and without implantation failure, Acta Obstet. Gynecol. Scand. 103 (2024) 1348–1365. 10.1111/aogs.14822.

[15] H. Heydarian, M. Abbasi, F. Najafi, M. Darbandi, Data Mining of Infertility and Factors Influencing Its Development: A Finding From a Prospective Cohort Study of RaNCD in Iran, Health Sci. Rep. 8 (2025) e70265. 10.1002/hsr2.70265.

[16] A.D.S. Pathare, K. Zaveri, I. Hinduja, Downregulation of genes related to immune and inflammatory response in IVF implantation failure cases under controlled ovarian stimulation, Am. J. Reprod. Immunol. N. Y. N 1989 78 (2017). 10.1111/aji.12679.

[17] N. Salem, S. Hussein, Data dimensional reduction and principal components analysis, Procedia Comput. Sci. 163 (2019) 292–299. 10.1016/j.procs.2019.12.111.

[18] Evans, C., Hardin, J., & Stoebel, D. M. (2018). Selecting between-sample RNA-Seq normalization methods from the perspective of their assumptions. Briefings in bioinformatics, 19(5), 776–792. 10.1093/bib/bbx008, (n.d.).

[19] M. Greenacre, P.J.F. Groenen, T. Hastie, A.I. D’Enza, A. Markos, E. Tuzhilina, Principal component analysis, Nat. Rev. Methods Primer 2 (2022) 100. 10.1038/s43586-022-00184-w.

[20] M.I. Love, W. Huber, S. Anders, Moderated estimation of fold change and dispersion for RNA-seq data with DESeq2, Genome Biol. 15 (2014) 550. 10.1186/s13059-014-0550-8.

[21] Benjamini, Y., & Hochberg, Y. (1995). Controlling the False Discovery Rate: A Practical and Powerful Approach to Multiple Testing. Journal of the Royal Statistical Society. Series B (Methodological), 57(1), 289–300. http://www.jstor.org/stable/2346101, (n.d.).

[22] C. Guangchuang Yu [Aut, clusterProfiler, (2017). 10.18129/B9.BIOC.CLUSTERPROFILER.

[23] M. Ashburner, C.A. Ball, J.A. Blake, D. Botstein, H. Butler, J.M. Cherry, A.P. Davis, K. Dolinski, S.S. Dwight, J.T. Eppig, M.A. Harris, D.P. Hill, L. Issel-Tarver, A. Kasarskis, S. Lewis, J.C. Matese, J.E. Richardson, M. Ringwald, G.M. Rubin, G. Sherlock, Gene Ontology: tool for the unification of biology, Nat. Genet. 25 (2000) 25–29. 10.1038/75556.

[24] M. Kanehisa, KEGG: Kyoto Encyclopedia of Genes and Genomes, Nucleic Acids Res. 28 (2000) 27–30. 10.1093/nar/28.1.27.

[25] Ingenuity Pathway Analysis (IPA), (n.d.). https://digitalinsights.qiagen.com/IPA.

[26] Krämer, A., Green, J., Pollard, J., Jr, & Tugendreich, S. (2014). Causal analysis approaches in Ingenuity Pathway Analysis. Bioinformatics (Oxford, England), 30(4), 523–530. 10.1093/bioinformatics/btt703, (n.d.).

[27] P. Langfelder, S. Horvath, WGCNA: an R package for weighted correlation network analysis, BMC Bioinformatics 9 (2008) 559. 10.1186/1471-2105-9-559.

[28] M. Pournasir, S. Ghorbian, T. Ghasemnejad, A. Fattahi, M. Nouri, Glutathione peroxidase 3 (extracellular isoform) levels and functional polymorphisms in fertile and infertile men, Middle East Fertil. Soc. J. 26 (2021) 13. 10.1186/s43043-021-00057-4.

[29] K. Palikaras, E. Lionaki, N. Tavernarakis, Balancing mitochondrial biogenesis and mitophagy to maintain energy metabolism homeostasis, Cell Death Differ. 22 (2015) 1399–1401. 10.1038/cdd.2015.86.

[30] M.S. Tomar, A. Kumar, A. Shrivastava, Mitochondrial metabolism as a dynamic regulatory hub to malignant transformation and anti-cancer drug resistance, Biochem. Biophys. Res. Commun. 694 (2024) 149382. 10.1016/j.bbrc.2023.149382.

[31] M. Idrees, Z. Haider, C.D. Perera, S. Ullah, S.-H. Lee, S.E. Lee, S.-S. Kang, S.W. Kim, I.-K. Kong, PPARGC1A regulates transcriptional control of mitochondrial biogenesis in early bovine embryos, Front. Cell Dev. Biol. 12 (2025) 1531378. 10.3389/fcell.2024.1531378.

[32] A. Fleig, R. Penner, The TRPM ion channel subfamily: molecular, biophysical and functional features, Trends Pharmacol. Sci. 25 (2004) 633–639. 10.1016/j.tips.2004.10.004.

[33] L.M. Pitzer, M.R. Moroney, N.J. Nokoff, M.J. Sikora, WNT4 Balances Development vs Disease in Gynecologic Tissues and Women’s Health, Endocrinology 162 (2021) bqab093. 10.1210/endocr/bqab093.

[34] Y.C. Ruan, H. Chen, H.C. Chan, Ion channels in the endometrium: regulation of endometrial receptivity and embryo implantation, Hum. Reprod. Update 20 (2014) 517–529. 10.1093/humupd/dmu006.

[35] L. Chen, D. Xiao, F. Tang, H. Gao, X. Li, CAPN6 in disease: An emerging therapeutic target (Review), Int. J. Mol. Med. (2020). 10.3892/ijmm.2020.4734.

[36] J. Mu, W. Wang, B. Chen, L. Wu, B. Li, X. Mao, Z. Zhang, J. Fu, Y. Kuang, X. Sun, Q. Li, L. Jin, L. He, Q. Sang, L. Wang, Mutations in *NLRP2* and *NLRP5* cause female infertility characterised by early embryonic arrest, J. Med. Genet. 56 (2019) 471–480. 10.1136/jmedgenet-2018-105936.

[37] P. Zhang, M. Dixon, M. Zucchelli, F. Hambiliki, L. Levkov, O. Hovatta, J. Kere, Expression Analysis of the NLRP Gene Family Suggests a Role in Human Preimplantation Development, PLoS ONE 3 (2008) e2755. 10.1371/journal.pone.0002755.

[38] D. Fotiadis, Y. Kanai, M. Palacín, The SLC3 and SLC7 families of amino acid transporters, Mol. Aspects Med. 34 (2013) 139–158. 10.1016/j.mam.2012.10.007.

[39] X. Hu, F. Chen, L. Jia, A. Long, Y. Peng, X. Li, J. Huang, X. Wei, X. Fang, Z. Gao, M. Zhang, X. Liu, Y.-G. Chen, Y. Wang, H. Zhang, Y. Wang, A gut-derived hormone regulates cholesterol metabolism, Cell 187 (2024) 1685–1700.e18. 10.1016/j.cell.2024.02.024.

[40] A. Fukui, M. Kamoi, A. Funamizu, K. Fuchinoue, H. Chiba, M. Yokota, R. Fukuhara, H. Mizunuma, NK cell abnormality and its treatment in women with reproductive failures such as recurrent pregnancy loss, implantation failures, preeclampsia, and pelvic endometriosis, Reprod. Med. Biol. 14 (2015) 151–157. 10.1007/s12522-015-0207-7.

[41] K. Sfakianoudis, A. Rapani, S. Grigoriadis, A. Pantou, E. Maziotis, G. Kokkini, C. Tsirligkani, S. Bolaris, K. Nikolettos, M. Chronopoulou, K. Pantos, M. Simopoulou, The Role of Uterine Natural Killer Cells on Recurrent Miscarriage and Recurrent Implantation Failure: From Pathophysiology to Treatment, Biomedicines 9 (2021) 1425. 10.3390/biomedicines9101425.

[42] S. Vandendriessche, S. Cambier, P. Proost, P.E. Marques, Complement Receptors and Their Role in Leukocyte Recruitment and Phagocytosis, Front. Cell Dev. Biol. 9 (2021) 624025. 10.3389/fcell.2021.624025.

[43] R. Bulla, F. Bossi, F. Tedesco, The Complement System at the Embryo Implantation Site: Friend or Foe?, Front. Immunol. 3 (2012). 10.3389/fimmu.2012.00055.

[44] J.E. Salmon, C. Heuser, M. Triebwasser, M.K. Liszewski, D. Kavanagh, L. Roumenina, D.W. Branch, T. Goodship, V. Fremeaux-Bacchi, J.P. Atkinson, Mutations in Complement Regulatory Proteins Predispose to Preeclampsia: A Genetic Analysis of the PROMISSE Cohort, PLoS Med. 8 (2011) e1001013. 10.1371/journal.pmed.1001013.

[45] S. Zhao, J. Lu, Y. Chen, Z. Wang, J. Cao, Y. Dong, Exploration of the potential roles of m6A regulators in the uterus in pregnancy and infertility, J. Reprod. Immunol. 146 (2021) 103341. 10.1016/j.jri.2021.103341.

[46] I.D. Keleş, T. Günel, B.Y. Özgör, E. Ülgen, E. Gümüşoğlu, M.K. Hosseini, U. Sezerman, F. Buyru, J. Yeh, E. Baştu, Gene pathway analysis of the endometrium at the start of the window of implantation in women with unexplained infertility and unexplained recurrent pregnancy loss: is unexplained recurrent pregnancy loss a subset of unexplained infertility?, Hum. Fertil. Camb. Engl. 26 (2023) 1129–1141. 10.1080/14647273.2022.2143299.

[47] A. Ferramosca, V. Zara, Diet and Male Fertility: The Impact of Nutrients and Antioxidants on Sperm Energetic Metabolism, Int. J. Mol. Sci. 23 (2022) 2542. 10.3390/ijms23052542.

[48] J. He, M. Ma, Z. Xu, J. Guo, H. Chen, X. Yang, P. Chen, G. Liu, Association between semen microbiome disorder and sperm DNA damage, Microbiol. Spectr. 12 (2024) e0075924. 10.1128/spectrum.00759-24.

[49] L. Zhang, Y. Feng, Y. Zhang, X. Sun, Q. Ma, F. Ma, The Sweet Relationship between the Endometrium and Protein Glycosylation, Biomolecules 14 (2024) 770. 10.3390/biom14070770.

[50] J. Hu, Z. Zhang, W.-J. Shen, S. Azhar, Cellular cholesterol delivery, intracellular processing and utilization for biosynthesis of steroid hormones, Nutr. Metab. 7 (2010) 47. 10.1186/1743-7075-7-47.

[51] Z. Zhou, J. Cao, X. Liu, M. Li, Evidence for the butyrate metabolism as key pathway improving ulcerative colitis in both pediatric and adult patients, Bioengineered 12 (2021) 8309–8324. 10.1080/21655979.2021.1985815.

[52] S. Gökben, S. Yılmaz, J. Klepper, G. Serdaroğlu, H. Tekgül, Video/EEG recording of myoclonic absences in GLUT1 deficiency syndrome with a hot-spot R126C mutation in the SLC2A1 gene, Epilepsy Behav. EB 21 (2011) 200–202. 10.1016/j.yebeh.2011.03.027.

[53] Y. Gong, G. Zhao, Wealth, health, and beyond: Is COVID-19 less likely to spread in rich neighborhoods?, PloS One 17 (2022) e0267487. 10.1371/journal.pone.0267487.

[54] S. Han, X. Zhang, J. Ding, X. Li, X. Zhang, X. Jiang, S. Duan, B. Sun, X. Hu, Y. Gao, Serum metabolic profiling of rats infected with Clonorchis sinensis using LC-MS/MS method, Front. Cell. Infect. Microbiol. 12 (2022) 1040330. 10.3389/fcimb.2022.1040330.

[55] C.L. Chang, W.-C. Lo, T.-H. Lee, J.-Y. Sung, Y.J. Sung, Oocyte-specific disruption of adrenomedullin 2 gene enhances ovarian follicle growth after superovulation, Front. Endocrinol. 13 (2022) 1047498. 10.3389/fendo.2022.1047498.

[56] N.K. Krishna, K.M. Cunnion, G.A. Parker, The EPICC Family of Anti-Inflammatory Peptides: Next Generation Peptides, Additional Mechanisms of Action, and In Vivo and Ex Vivo Efficacy, Front. Immunol. 13 (2022) 752315. 10.3389/fimmu.2022.752315.

[57] O. Wijaya, J.Y. Anas, W. Widjiati, M.Y.A. Widyanugraha, S. Samsulhadi, H. Bayuaji, S.R. Dwiningsih, B.Y. Utomo, B. Stevanny, Altered Mitochondrial Morphology and Reduced Cardiolipin Levels in Oocytes of Endometriosis Model Mice: Implications for Mitochondrial Dysfunction in Infertility, Med. Sci. Monit. Int. Med. J. Exp. Clin. Res. 31 (2025) e947194. 10.12659/MSM.947194.

[58] C. Mette, B. Camilla Dooleweerdt, J. Stine, B. Anders Miki, P. Morten Roenn, L.-J. Henrik, Evaluation of the systemic acute phase response and endometrial gene expression of serum amyloid A and pro- and anti-inflammatory cytokines in mares with experimentally induced endometritis, Vet. Immunol. Immunopathol. 138 (2010) 95–105. 10.1016/j.vetimm.2010.07.011.

[59] S. Dallel, M. Despalles, M. Tore, Y. Renaud, A. Kocer, C. Damon-Soubeyrand, P. Pouchin, C. Vachias, K. Boutourlinsky, C. Gonthier-Gueret, A. De Haze, P. Sanchez, J.-C. Pointud, E. Bouchareb, M. Vialat, A. Lagarde, C. Gulunga, L. Chaput, A. Vega, F. Brugnon, I. Tauveron, A. Trousson, C. De Joussineau, F. Degoul, L. Morel, J.M. Lobaccaro, S. Maqdasy, S. Baron, LXR pathway drives hormonal response intensity in polycystic ovary syndrome, EMBO Mol. Med. 17 (2025) 1666–1685. 10.1038/s44321-025-00251-1.

[60] M.L. McCallum, C.A. Pru, A.R. Smith, N.C. Kelp, M. Foretz, B. Viollet, M. Du, J.K. Pru, A functional role for AMPK in female fertility and endometrial regeneration, Reproduction 156 (2018) 501–513. 10.1530/REP-18-0372.

[61] I. Tamura, Y. Doi□Tanaka, A. Takasaki, K. Shimamura, T. Yoneda, H. Takasaki, A. Shiroshita, T. Fujimura, Y. Shirafuta, N. Sugino, High incidence of decidualization failure in infertile women, Reprod. Med. Biol. 23 (2024) e12580. 10.1002/rmb2.12580.

[62] J.E. Thaxton, S. Sharma, REVIEW ARTICLE: Interleukin□10: A Multi□Faceted Agent of Pregnancy, Am. J. Reprod. Immunol. 63 (2010) 482–491. 10.1111/j.1600-0897.2010.00810.x.

[63] S. Seshadri, S.K. Sunkara, Natural killer cells in female infertility and recurrent miscarriage: a systematic review and meta-analysis, Hum. Reprod. Update 20 (2014) 429–438. 10.1093/humupd/dmt056.

[64] C. Uzuner, J. Mak, F. El-Assaad, G. Condous, The bidirectional relationship between endometriosis and microbiome, Front. Endocrinol. 14 (2023) 1110824. 10.3389/fendo.2023.1110824.

[65] C. Guo, C. Zhang, Role of the gut microbiota in the pathogenesis of endometriosis: a review, Front. Microbiol. 15 (2024) 1363455. 10.3389/fmicb.2024.1363455.

[66] I. Fernández-Ruiz, A newly identified gut hormone suppresses cholesterol production in the liver, Nat. Rev. Cardiol. 21 (2024) 358–358. 10.1038/s41569-024-01021-1.

