## Supplementary material for "Transcriptomic Analysis Reveals Inflammatory and Metabolic Dysregulation in Unexplained Female Infertility": https://docs.google.com/document/d/1GI6tYFj_wQyU3uO2r44K2myeRFkQ8hdQ/edit?usp=drive_link&ouid=100845841708565570348&rtpof=true&sd=true

Table 1: Significantly Upregulated KEGG Pathways

| **Pathway** | **p-value** | **Key Genes** | **Reference** |
| --- | --- | --- | --- |
| **Hematopoietic Cell Lineage** | **0.0005** | **CR1, CSF1R, TNF, CD3E** | **PMID: 34105167** |
| **Natural Killer Cell Mediated Cytotoxicity** | **0.0008** | **PIK3 family, TNF** | **PMID: 34116483** |
| **Complement and Coagulation Cascades** | **0.0014** | **CR1, TNF** | **PMID: 36369952** |
| **Chemokine Signaling Pathway** | **0.0017** | **PIK3CB, PIK3CD, TNF** | **PMID: 33936247** |
| **Antigen Processing and Presentation** | **0.0120** | **PIK3CD, PIK3CB** | **PMID: 38201263** |
| **Fc epsilon RI Signaling Pathway** | **0.0291** | **PIK3CA, PIK3CB** | **PMC ID: Mast Cell-Sperm Function** |

**Table 2: Significantly Downregulated KEGG Pathways**

| **Pathway** | **p-value** | **Key Genes** | **Reference** |
| --- | --- | --- | --- |
| **Terpenoid Backbone Biosynthesis** | **0.0148** | **PMVK, FDPS, HMGCR, HMGCS1, ACAT1, ACAT2** | **PMID: 32923184** |
| **Biosynthesis of Unsaturated Fatty Acids** | **0.0245** | **HADHA, FADS1, ELOVL2, SCD, SCD5** | **PMID: 35269682** |
| **Butanoate Metabolism** | **0.0260** | **HADHA, HMGCL, ACAT1, ACAT2** | **PMID: 38899893** |
| **Propanoate Metabolism** | **0.0290** | **ECHS1, ALDH2, ACAT1, ACAT2, PDHB** | **PMID: 40220430** |
| **Mucin Type O-Glycan Biosynthesis** | **0.0377** | **GALNT6, GALNT13, GCNT1, ST6GALNAC1** | **PMID: 39062484** |
| **Steroid Biosynthesis** | **0.0455** | **CYP2R1, CYP27B1, CYP51A1, HSD17B7** | **PMID: 35269682** |

We found 3 activated and 5 significantly inhibited canonical pathways Table (1).

| **Ingenuity Canonical Pathways** | **-log(p-value)** | **z-score** | **Molecules** |
| --- | --- | --- | --- |
| Acute Phase Response Signaling | 3.40E+00 | 2.236 | C3,C4BPA,FGA,FGB,IL6ST,ORM1,ORM2,SOD2 |
| LXR/RXR Activation | 2.15E+00 | 2.236 | C3,FGA,ORM1,ORM2,PTGS2 |
| Response to elevated platelet cytosolic Ca2+ | 1.47E+00 | 2 | FGA,FGB,ORM1,ORM2 |
| 3-phosphoinositide Biosynthesis | 1.83E+00 | -2 | ACP6,ALPL,PIK3C2G,PIP5K1B,PPP1R14C,PSPH |
| Adrenomedullin signaling pathway | 1.41E+00 | -2 | C3,GAD1,GPR37,PIK3C2G,PLCH1 |
| Ion channel transport | 3.52E+00 | -2.121 | ATP12A,ATP6V0E2,ATP8A1,ATP8B3,CAMK2B,TRPM6,TTYH2,WNK4 |
| Super pathway of Inositol Phosphate Compounds | 2.15E+00 | -2.236 | ACP6,ALPL,PIK3C2G,PIP5K1B,PLCH1,PPP1R14C,PSPH |
| Mitochondrial Dysfunction | 2.32E+00 | -2.333 | ATP12A,CAMK2B,CAPN6,CREB3L1,GPX3,HAP1,PIK3C2G,PPARGC1A,SOD2 |

| Diseases or Functions Annotation | p-value | Predicted Activation State | Activation z-score |
| --- | --- | --- | --- |
| Obesity | 3.93E-03 | Increased | 2.792 |
| Inflammation of gastrointestinal tract | 4.29E-07 | Increased | 2.736 |
| Gastroenteritis | 4.16E-06 | Increased | 2.736 |
| Enteritis | 1.52E-05 | Increased | 2.668 |
| Abnormality of large intestine | 6.06E-06 | Increased | 2.484 |
| Colitis | 2.91E-05 | Increased | 2.289 |
| Quantity of carbohydrate | 3.99E-05 | Increased | 2.256 |
| Concentration of D-glucose | 1.08E-04 | Increased | 2.234 |
| Weight loss | 1.40E-03 | Increased | 2.164 |
| Cell movement of sperm | 7.75E-03 | Decreased | -2 |
| Transport of ion | 6.51E-07 | Decreased | -2.138 |
| Development of head | 1.15E-03 | Decreased | -2.415 |
| Transport of monovalent inorganic cation | 7.03E-05 | Decreased | -2.433 |
